# Development of natural scene representation in primary visual cortex requires early postnatal experience

**DOI:** 10.1101/2020.10.14.338897

**Authors:** Nina Kowalewski, Janne Kauttonen, Patricia L. Stan, Brian B. Jeon, Thomas Fuchs, Steven M. Chase, Tai Sing Lee, Sandra J. Kuhlman

**Affiliations:** Department of Biological Sciences, Carnegie Mellon University, 4400 Fifth Ave, Pittsburgh, PA 15213, USA; Center for the Neural Basis of Cognition, 1400 Locust Street, Pittsburgh, PA 15219, USA; University of Pittsburgh Center for Neuroscience, 4400 Fifth Ave, Pittsburgh, PA 15213, USA; Department of Biomedical Engineering, Carnegie Mellon University, 4400 Fifth Ave, Pittsburgh, PA 15213, USA; Neuroscience Institute, Carnegie Mellon University, 4400 Fifth Ave, Pittsburgh, PA 15213, USA; Department of Computer Science, Carnegie Mellon University, 4400 Fifth Ave, Pittsburgh, PA 15213, USA

**Author notes:** co-first authors (Nina Kowalewski, Janne Kauttonen). Correspondence should be addressed to: Sandra Kuhlman,.

## Abstract

The development of the visual system is known to be shaped by early-life experience. To identify response properties that contribute to enhanced natural scene representation, we performed calcium imaging of excitatory neurons in the primary visual cortex (V1) of awake mice raised in three different conditions (standard-reared, dark-reared, and delayed-visual experience) and compared neuronal responses to natural scene features relative to simpler grating stimuli that varied in orientation and spatial frequency. We assessed population selectivity in V1 using decoding methods and found that natural scene discriminability increased by 75% between the ages of 4 to 6 weeks. Both natural scene and grating discriminability were higher in standard-reared animals compared to those raised in the dark. This increase in discriminability was accompanied by a reduction in the number of neurons that responded to low-spatial frequency gratings. At the same time there was an increase in neuronal preference for natural scenes. Light exposure restricted to a 2-4 week window during adulthood did not induce improvements in natural scene nor in grating stimulus discriminability. Our results demonstrate that experience reduces the number of neurons required to effectively encode grating stimuli and that early visual experience enhances natural scene discriminability by directly increasing responsiveness to natural scene features.

## Introduction

During early postnatal development, sensory systems have the potential to adapt to complex natural scene patterns in the local environment in many ways. First, it has been observed that visual experience refines neural responses to simple grating stimuli such that response properties are matched to the local scene statistics experienced by the animal [1–8]. These observations raise the possibility that refinement of responses to low level features is sufficient to improve natural scene discriminability [9,10]. For example, it has been proposed that experience regulates orientation tuning diversity at the population level to match the environment and that natural scenes are effectively encoded by a synergistic interaction between a distribution of neurons broadly and narrowly tuned for orientation [11].

However, it has also long been recognized that some neurons within the primary visual cortex (V1) respond specifically to complex stimuli, including natural scenes [12–17]. In addition, it is becoming increasingly apparent that selectivity to complex features found in natural scenes cannot be explained by response profiles evoked by simple features such as grating stimuli [18–23], and that this is likely a general principle of sensory processing [24]. Selectivity to natural scenes at the level of V1 is functionally significant, as it can account for accuracy of scene discrimination in primates [25]. These studies raise the possibility that experience drives individual V1 neurons to become directly sensitive to complex features found in natural scenes. Complex features include higher order statistics and spatial correlations [17,26,27], multi-edged elements and non-Cartesian shapes within the classic receptive field [14,21,28], as well as elements and shapes extending beyond the classical receptive field [19,29,30]. Tuning to these higher order features will likely lead to greater representational efficiency and better discrimination of complex natural scene stimuli.

The goal of this study was to evaluate the impact of experience during development on the responsiveness and preference of V1 neurons in the mouse to natural scene stimuli and assess how experience-induced changes in natural scene selectivity are related to classic orientation and spatial frequency tuning properties. Two-photon calcium imaging was used to image the neural responses of hundreds of neurons simultaneously in awake head-fixed mice raised in either standard-reared (SR), dark-reared (DR), or delayed visual experience (DVE) conditions. SR mice were housed continuously in standard lighting conditions on a 12 hour lights on/ off cycle. DR mice were placed in a light-tight dark cabinet from postnatal day (P) 6 until the day that they were first imaged. DVE-raised mice experienced vision as adults, after the age of P35, for approximately 2-4 weeks in standard lighting conditions. These mice did not experience vision prior to the age of P35. In a single imaging session, static natural scenes that varied in pairwise similarity as well as static gratings that included a wide range of spatial frequencies were presented. Natural scene discriminability of V1 neuronal populations in animals from these three rearing conditions were assessed based on stimulus decoding and in relation to other tuning properties, including orientation selectivity and spatial frequency preference. We found that the number of V1 neurons with a strong preference for natural scene stimuli was markedly higher in standard-reared animals compared to dark-reared animals. This difference was accompanied by a decrease in responsiveness to grating stimuli in standard-reared animals. The decrease was specific to the low spatial frequency range; higher spatial frequencies were not impacted by rearing condition. Delayed visual experience for 2-4 weeks in DVE adult mice was insufficient for neurons to fully develop the preference for natural scenes found in standard-reared animals. Finally, we found that preference for natural scene stimuli over grating stimuli was predictive of natural scene decoding performance on an animal-by-animal basis. Our results suggest that experience-dependent development of natural scene discriminability in V1 is not simply due to the refinement of low level spatial frequency or orientation tuning, but requires the development of new sensitivity to more complex natural scene features.

## Results

To assess the impact of visual experience on natural scene discriminability we selected a set of 20 natural scene images that varied in similarity by choosing individual image frames from two different movie sequences, a forest and a city sequence. Ten forest and ten city scenes were included. Activity of individual layer 2/3 V1 neurons was imaged in response to presentations of grating and natural scene images using 2-photon microscopy in head-fixed transgenic mice expressing the calcium indicator GCaMP6f (**Fig. 1A,B**). We confirmed that the natural scene (NS) stimulus set used in this study contained images that were easy as well as difficult for mice to perceptually discriminate in a Go/No-go behavioral task (**Figs. 1C, S1A**). Similarity of natural scene pairs was quantified using the structural similarity index (SSIM) [31]. Scene pairs with a high index are similar and are more difficult to discriminate compared to a scene pair with a lower index. The SSIM index was previously shown to be a reasonable indicator of perceptual discriminability [32], and in our case we used this index to confirm that frames close together in the movie sequence were more similar than distal frames (**Fig. S1B**). For the scenes presented, we found that mice were able to easily detect the difference between a scene pair having an SSIM of 0.67 or less, but exhibited lower performance for scene pairs with higher SSIM values. We considered pairwise comparisons of frames from the same sequence, forest or city, to be ‘within scene-type’, and pairwise comparisons between the forest and city sequences to be ‘cross scene-type’ comparisons. As expected, the power spectrum for both forest and city scenes contained the highest power in low spatial frequencies known to drive mouse V1 neurons, and also included spatial frequencies higher than what mice are sensitive to (**Fig. S1C**).

**Figure 1.**
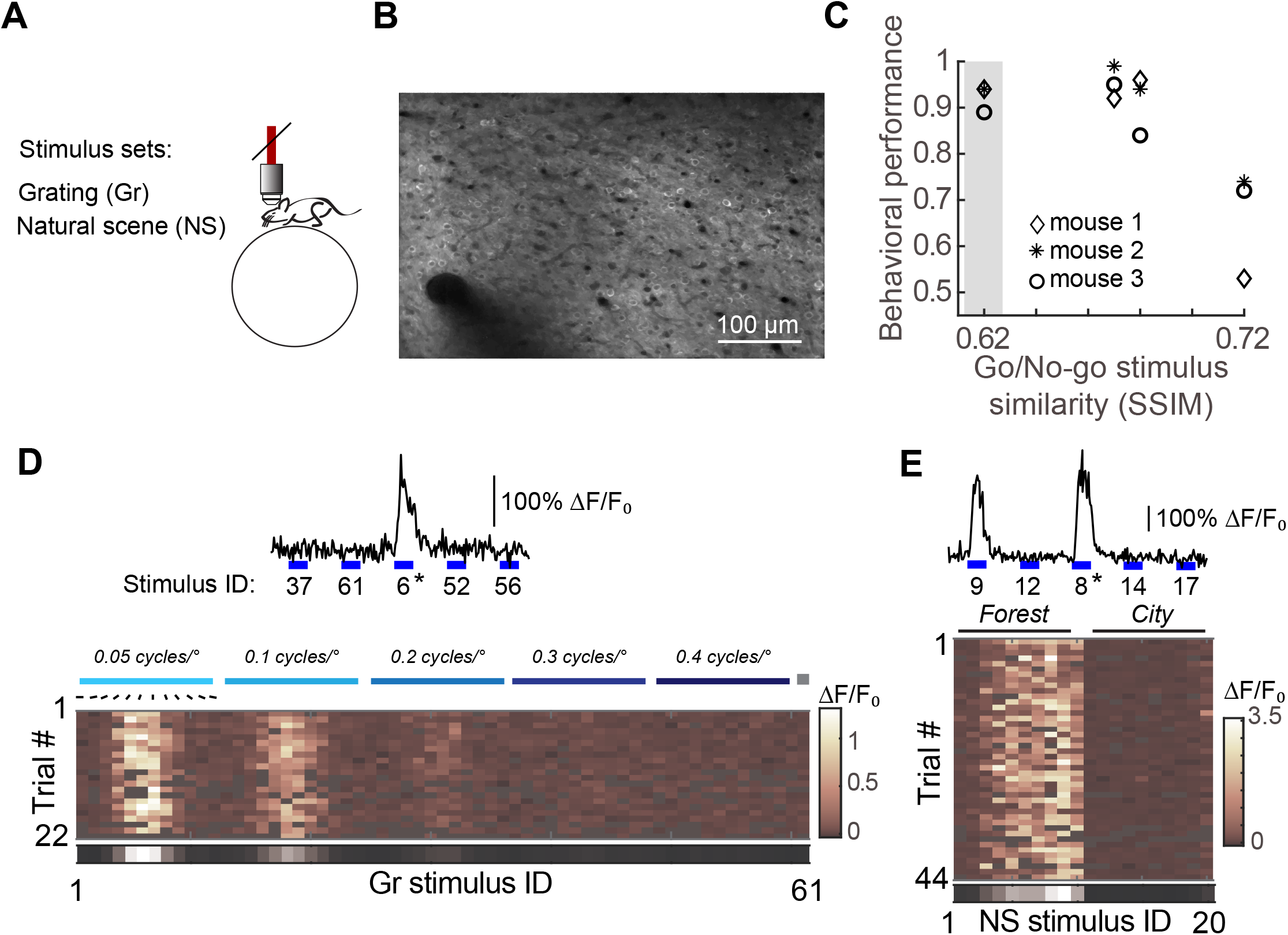
V1 neural responses to static natural images of varying similarity and grating stimuli. **A** Neural responses to grating and natural scene stimuli were recorded using 2-photon calcium imaging in awake mice, head-fixed atop a spherical treadmill. **B** Mean intensity 2-photon image averaged across 2,000 frames. A blood vessel is visible in lower left. Scale bar, 100 μm. **C** Go/No-go discrimination task performance decreased with increasing image similarity. Mice were shaped on a stimulus pair of low similarity (0.62 SSIM, gray shading), and tested on 3 additional stimulus pairs of varying difficulty (SSIM values: 0.67,0.68, 0.72). See **Fig. S1A** for further details on hit and false alarm rate. **D** Responses to grating stimuli, one example neuron. Stimulus trials were interleaved with a 2-second duration gray screen. Spatial frequencies as high as 0.4 cycles/° were tested at an orientation spacing of 15°. Top, continuous fluorescent trace, stimuli were randomized as indicated; a single preferred stimulus presentation is (*) shown flanked by two additional stimulus presentations. Stimulus duration (blue line) was 1 second, scale bar is 100% ΔF/F_0_. Bottom, signal amplitude was averaged across the full stimulus duration, individual stimulus trials were sorted as indicated. Stimulus ID # 61 was an additional blank gray screen (gray square), 1 second in duration. Trials removed due to motion artifact, locomotion, or eye movement are shown in gray, response amplitudes are scaled in color. The mean across trials is shown in gray scale at the bottom (scale limits: 0.7, 0 ΔF/F_0_). **F** Responses to 20 natural scene stimuli, one example neuron. Stimuli were interleaved with a 2-second duration gray screen, organized as in ‘D’.

In the same imaging session grating (Gr) stimuli of varying spatial frequency and orientation were presented so that responses to both natural scenes and gratings were captured for a given neuron. In total, 80 stimuli were presented. The highest grating spatial frequency presented was 0.4 cycles/°, the lowest was 0.05 cycles/°. In a separate experiment we confirmed that the range of spatial frequencies presented was close to saturating in terms of the number of neurons recruited. We found that 97± 0.01% of the neurons responding to spatial frequencies higher than 0.4 cycles/° also responded to at least one stimulus of 0.4 cycles/° or less (**Fig. S1D**). Grating stimuli and natural scene images were randomized and presented to the mice as static images interleaved with isoluminant gray screen presentations (**Fig. 1D,E**). Approximately 44 repeated trials per stimulus of natural scene images and approximately 22 repeated trials per stimulus of grating stimuli were presented, separated into blocks of roughly 10 and 5 consecutive trials, respectively. Using this presentation sequence, we saw no evidence of response adaptation within an imaging session (**Fig. S2A**). Locomotion as well as pupil position was monitored during imaging. Trials containing locomotion or eye movements greater than 10° from the median (**Fig. S2B-C**) were removed from further analysis.

### Impact of experience on natural scene discrimination

The influence of visual experience on V1 neural activity was first assessed by comparing the number of neurons that responded to grating stimuli in standard-reared versus dark-reared mice. Animals were raised in the dark (DR) until at least age P35 (9 animals) and compared with animals raised in standard conditions (SR, 8 animals). An advantage of calcium imaging compared to other recording techniques is that the total number of neurons sampled is a known quantity, therefore the proportion of responsive neurons can be calculated and compared between rearing conditions. We found that 66% of the imaged neurons from dark-reared animals responded to at least one of the 80 stimuli presented (2729 out of 4154 imaged neurons pooled across animals, 66%). The other 34% did not respond to any of the stimuli presented. The proportion of responsive neurons was similar in standard-reared animals (2552 out of 3819 imaged neurons responded, 67%; **Fig. 2A**).

**Figure 2.**
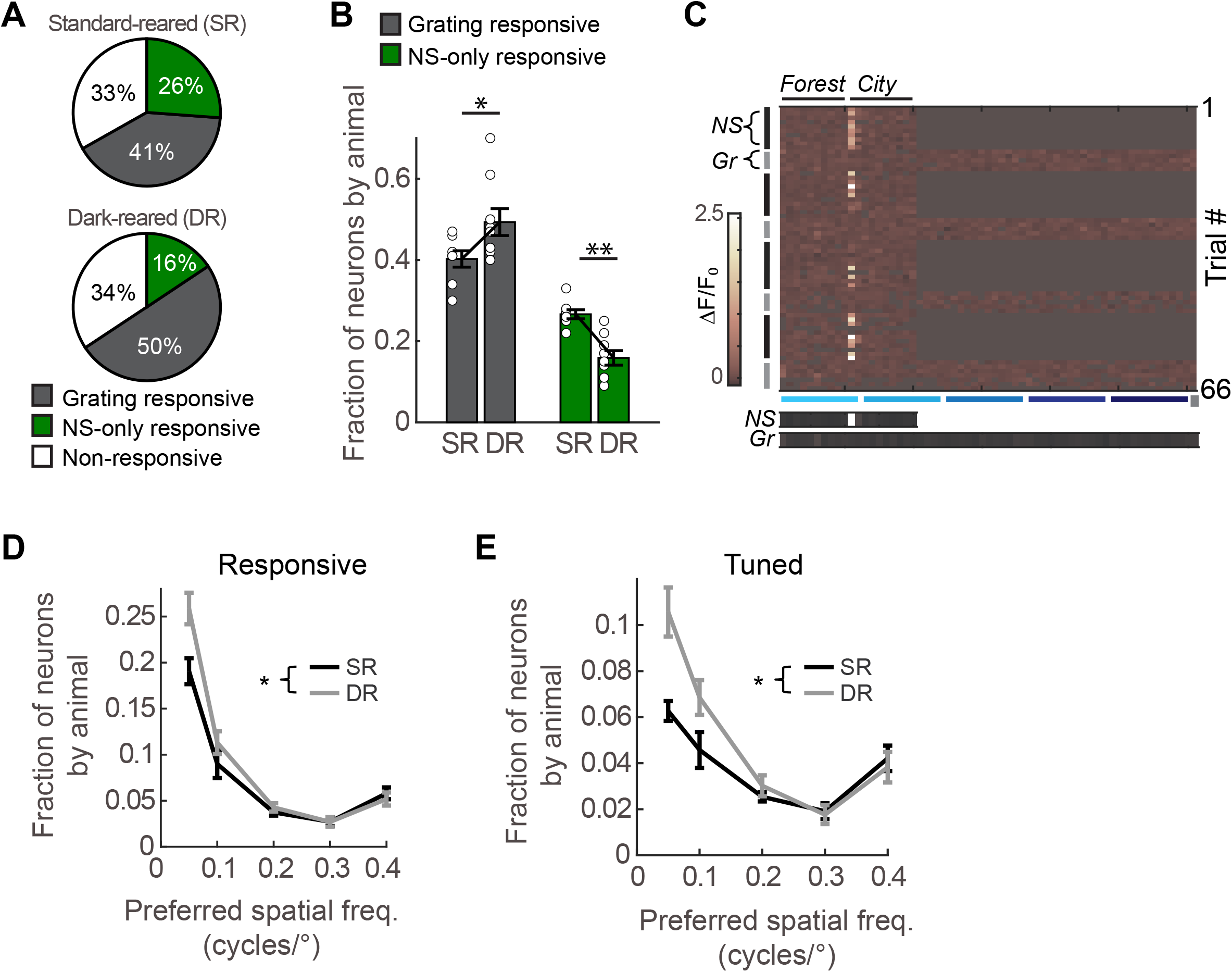
The number of grating-responsive neurons is higher in dark-reared mice. **A** The proportion of neurons responding to visual stimuli was similar in standard-reared (SR, 3819 total neurons from 8 animals) compared to dark-reared (DR, 4154 total neurons from 9 animals) animals. The grating-responsive category includes neurons that respond to both grating and natural scene stimuli. In the SR condition, 60% of the grating responsive neurons also responded to at least one natural scene, and in the DR condition 64% responded to at least one natural scene. **B** The number of neurons responding to grating stimuli was significantly higher in DR animals compared to SR animals (Wilcoxon rank test, p= 0.025). The number of neurons responding to natural scenes (NS) only was significantly lower in DR animals compared to SR animals (Wilcoxon rank test, p= 5.76e-4). Circles indicate individual animals (DR, n= 9 animals; SR, n=8 animals); the mean is indicated by the bar. **C** Responses to grating and natural scene stimuli, one example neuron that responded to only natural scenes. Natural Scene (NS, black vertical bar) and Grating stimuli (Gr, gray vertical bar) were presented in alternating blocks as indicated. The mean across NS and Gr trials is shown in gray scale at the bottom (scale limits: 0.7, 0 ΔF/F_0_). Stimulus IDs sorted as in Figure 1D,E. **D** Distribution of spatial frequency preference in SR and DR animals, all grating responsive neurons are included. The number of grating responsive neurons was significantly higher in DR conditions compared to SR and the difference between rearing conditions depended on spatial frequency (two-way mixed repeated measures ANOVA, between-subjects rearing condition p = 0.044; interaction between rearing condition and preferred spatial frequency p= 0.012 Greenhouse-Geisser corrected for sphericity). **E** Distribution of spatial frequency preference in SR and DR animals, only tuned neurons that were well-described by a 2-dimensional Gaussian fit are included. The number of grating responsive neurons was significantly higher in DR conditions compared to SR and the difference between rearing conditions depended on spatial frequency (two-way mixed repeated measures ANOVA, between-subjects rearing condition p = 0.038; interaction between rearing condition and preferred spatial frequency p= 0.001Greenhouse-Geisser corrected for sphericity). See **Fig. S3** for example neurons well-fit by a 2-dimensional Gaussian function.

Although the total number of responsive neurons was comparable, responsiveness to stimulus complexity was distinct between the two rearing conditions. First we considered the number of neurons that responded to simple grating stimuli. Neurons were included regardless of whether their grating stimulus responses were classically tuned to orientation and well fit by a Gaussian function. We found that the number of neurons that responded to at least one grating stimuli was higher in the dark reared condition. In dark-reared animals, half of the imaged neurons responded to grating stimuli, approximately 10% more than in standard reared animals (**Fig. 2A**). The higher number of grating responsive neurons relative to the standard reared condition was also evident on an animal-by animal basis (**Fig. 2B**). The higher number of grating-responsive neurons was accompanied by a 10% reduction in the number of neurons responding exclusively to one or more natural scene images (NS-only responsive). An example of a neuron responding to natural scenes but not grating stimuli is shown in Figure 2C. Further analysis revealed that the difference in grating responsiveness between the two rearing conditions was specific to low spatial frequencies. This effect was present regardless of whether all responsive neurons were considered or just those neurons that were classically tuned to orientation and spatial frequency (**Figs. 2D,E**). We did not test spatial frequencies below the most common preferred spatial frequency [33]. Given this, any neurons that respond to ≥0.05 cycles/° and have a spatial frequency preference less than 0.05 cycles/° are scored as having a spatial frequency of 0.05 cycles/° or higher. Therefore, it is possible that the proportion of neurons for each spatial frequency reported here are slightly over estimated, particularly for the lowest spatial frequency tested.

Notably, visual experience did not increase the proportion of total imaged neurons that responded to high spatial frequency grating stimuli. The proportion of neurons responsive to high frequency gratings was similar between the two rearing conditions. However, the relative distribution of neurons tuned to orientation and spatial frequency was shifted towards higher spatial frequencies. In other words, the U-shaped curve flattened on the left side. These results indicate visual experience drives V1 to become less responsive to low spatial frequency grating stimuli and become more responsive to natural scenes.

The above response properties are an indication that experience enhances the representation of natural scenes within V1. To directly test this, the ability of V1 neurons to discriminate the natural scene stimulus set was assessed using decoding methods. The data were analyzed on an animal-by animal basis using a nearest-neighbor decoder. We found that decoding accuracy was significantly higher in standard-reared animals compared to dark-reared animals (**Fig. 3A**). Similar results were obtained using a Naive Bayes decoder (**Fig. S4**), establishing that the observed effect was robust to the choice of decoder. Analysis of the confusion matrices, which include all 20 natural scene predictions (**Fig. 3B**), gave us an opportunity to determine which scene type comparisons (e.g. cross scene type which are dissimilar versus within scene-type which are similar) generated errors. We found that the probability of error for both cross scene-type and within scene-type was lower in standard-reared animals compared to dark-reared animals (**Fig. 3C,D**).

**Figure 3.**
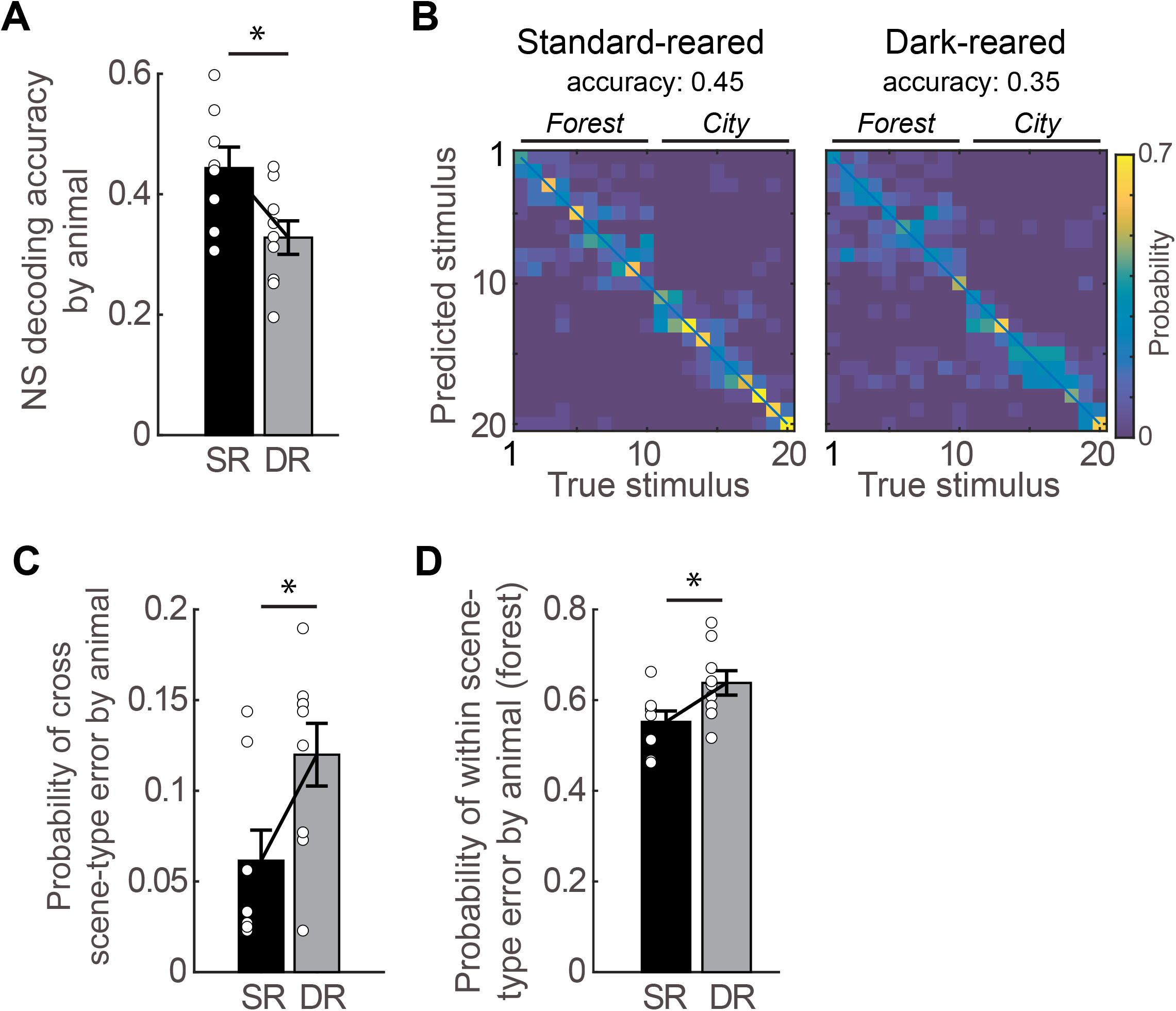
Natural scene discriminability in V1 is higher in standard-reared mice. **A** Decoding accuracy of natural scene (NS) stimuli was significantly lower in DR animals (n=9 animals) compared to SR animals (n= 8 animals; t-test, p= 0.019). 151 neurons per animal were used, 24 trials for each natural scene stimulus were included. Chance performance is 0.05. **B** Example confusion matrix plots from a SR animal and a DR animal. Accuracy is calculated as the sum of the diagonal (blue line). **C** Probability of cross-scene error (city and forest scene values were averaged) was significantly higher in DR compared to SR animals (Wilcoxon rank test, p= 0.034). **D** Probability of within-scene error, specifically for the forest scene type, was significantly higher in DR compared to SR animals (Wilcoxon rank test, p= 0.036). Error bars indicate S.E.M. *p<0.05, **p<0.01

In addition to sensitivity, we also examined whether stimulus preference was distinct between the rearing conditions. We found that many visually responsive neurons responded strongly to both natural scene and grating stimuli (**Fig. 4A**), however the majority of neurons responded strongly to either natural scene stimuli (**Fig. 2C**) or grating stimuli (**Fig. 4A**). Of the neurons that responded to grating stimuli, some were classically tuned (**Fig. 4A, middle**), and were well-described by a 2-dimensional Gaussian fit (**Fig. S3**). We also observed grating responsive neurons that did not exhibit classic tuning characteristics (**Fig. 4A, right**) [34,35]. In this later example, the neuron selectively responded to a subset or orientations for the lowest spatial frequency presented as well as for high spatial frequencies, but not to spatial frequencies of 0.1 cycles/°, similar to a previous report [34]. We quantified the relative preference for grating vs natural scene stimuli on a neuron-by-neuron basis as the natural scene-grating (NG) preference index. The NG preference index does not take into account tuning properties; the only criteria for this analysis is that the neuron responds to at least one stimulus from either stimulus set (grating or natural scene), so neurons such as those shown in Figure 4A right are included. Neurons responding robustly to both grating and natural scenes comprised 36.6% of all the imaged neurons, pooled across animals. The median NG preference index of all responsive neurons pooled across standard-reared animals was 0.17, and was shifted rightward compared to dark-reared animals which had a mean NG preference index of −0.20 (**Fig. 4B**). The difference between rearing conditions was also evident on an animal-by-animal basis in which the median NG preference index of the neurons within an animal was calculated, and compared across rearing conditions, the average difference between SR and DR conditions was 0.45 (**Fig. 4C**).

**Figure 4.**
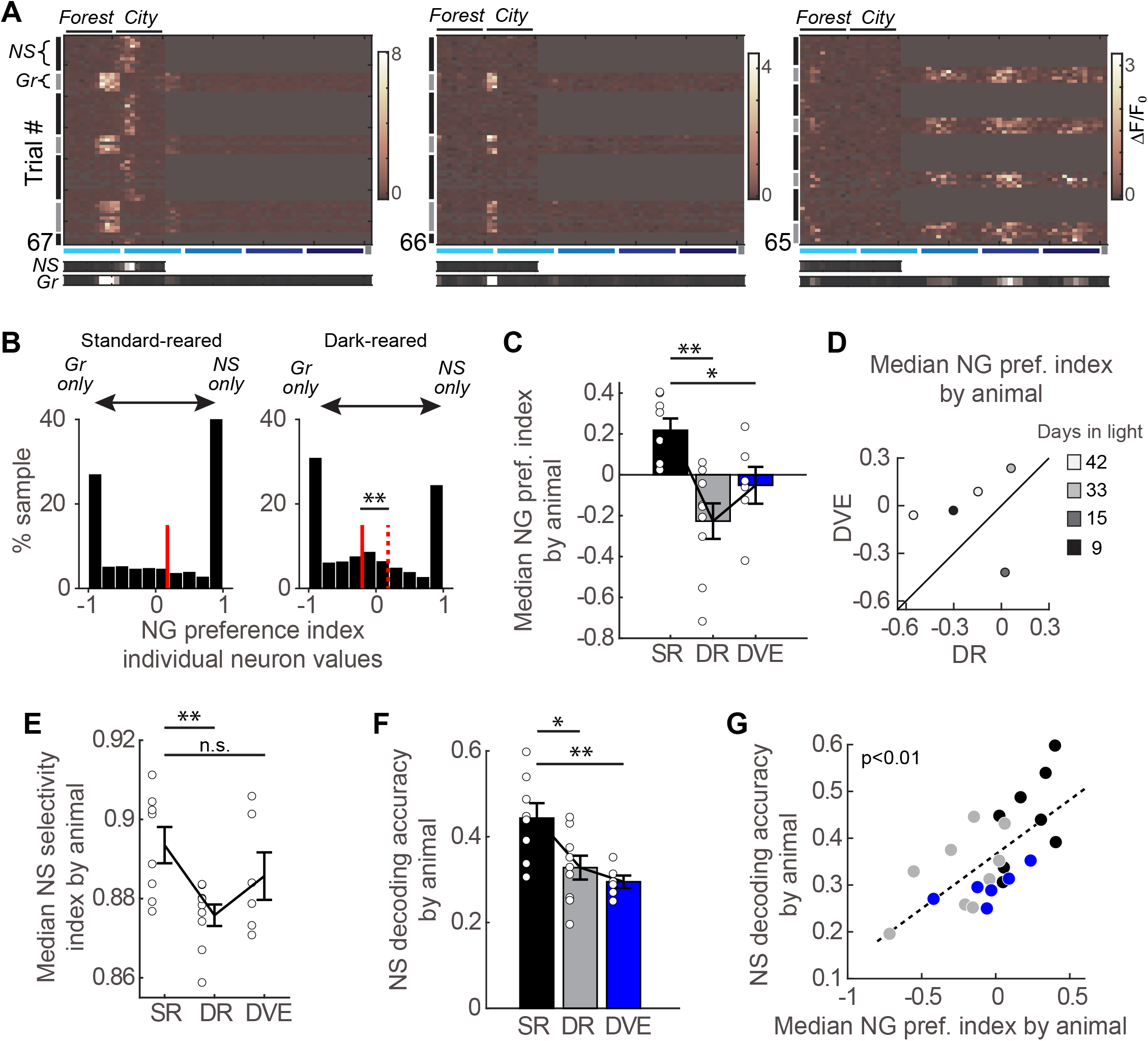
The proportion of neurons preferring natural scenes over grating stimuli is predictive of discrimination accuracy in V1. **A** Three example neuron responses to natural scenes (*NS*) and grating stimuli (*Gr*), organized as in Fig. 2C. **B** NG preference index distribution of individual neuron responses is significantly different between the SR (2551 neurons pooled from 8 standard-reared animals) and DR (2726 neurons pooled form 9 dark-reared animals) rearing conditions (KS-test, p= 3.35e-30). Median indicated in red, dashed line is the median of the SR condition. **C** Median NG preference index for each animal (circles) and the average of median values (bar). A value of 1 indicates the neuron did not respond to any of the grating stimuli (NS only), and a value of −1 indicates the neuron did not respond to any of the natural scene stimuli (GR only). See **Fig. S5** for example neurons. The difference between SR (n= 8 animals) and DR (n= 9 animals) as well as SR and DVE (n= 6 animals) was significant (t-test, Bonferroni-corrected for 2 planned comparisons, p=0.002 and p=0.043 respectively). **D** Five animals were successfully transitioned from DR to DVE. Four of the five animals has a positive shift in NG preference. Line indicates unity. **E** Median natural scene (NS) selectivity index for each animal (circles) by rearing condition and the average of median values (bar). The difference between SR (n= 8 animals) and DR (n= 9 animals) but not SR and DVE (n= 6 animals) was significant (t-test, Bonferroni-corrected for 2 comparisons, p=0.008 and p=0.526 respectively). **F** Decoding accuracy of natural scene (NS) stimuli was significantly lower in DVE animals (n=6 animals) compared to SR animals (n= 8 animals; t-test, Bonferroni-corrected for 2 planned comparisons, p= 0.009). 151 neurons per animal were used, 24 trials for each natural scene stimulus were included. SR and DR data are re-plotted from **Fig. 2F**. **G** Natural scene decoding accuracy correlated with the NG selectivity index on an animal-by-animal basis. Pearson correlation coefficient r= 0.67, p= 4.94e-4. Data points are colored by condition, SR: black, DR: gray, DVE: blue. Error bars indicate S.E.M. *p<0.05, **p<0.01

We were able to transition a subset of the dark-reared animals to standard conditions. Six dark-reared mice were placed in standard lighting conditions immediately after their first imaging session, which occurred at age p35 or older, and were then imaged approximately 32 days later (median days in standard conditions was 32, the range was 9 - 42 days). This condition is referred to as delayed visual experience (DVE). The median NG preference index in DVE animals was significantly lower compared to standard-reared animals (**Fig. 4C**). This is an indication that 30 days of visual experience as an adult is insufficient to fully shift the preference towards natural scenes to the amount seen in normal reared animals. Consistent with this interpretation, for those animals that were imaged both in the DR and DVE conditions, in 4 out of 5 animals the difference between the two conditions was less than 0.45, the difference between SR and DR conditions (**Fig.4D**). In the one animal that the shift was greater than 0.45, the absolute value after 42 days in light (−0.06) was well below the average value for the SR condition (0.22).

Next we considered response selectivity to just natural scenes. To quantify the response selectivity of individual neurons, we expressed natural scene (NS) selectivity as an index ranging from 0 to 1. Larger values indicate higher selectivity. Only neurons that responded to at least one natural scene and had a NS selectivity SEM < 0.1 were included. To compare rearing conditions, the median NS selectivity index of the neurons within an animal was computed. We found that natural scene selectivity to the 20 presented scenes was significantly lower in dark-reared animals compared to standard-reared animals, and that in contrast to the NG preference index, DVE animals were not different from standard-reared animals (**Fig. 4E**). Thus, early-life experience as well as adult experience later in life was sufficient to sharpen natural scene selectivity.

Is the observed increase in NS selectivity induced by late-life experience sufficient to improve discriminability to standard-reared levels? To address this, we compared decoding accuracy between standard-reared and DVE-reared animals. Decoding accuracy was significantly lower in DVE-raised animals compared to standard-reared animals (**Fig. 4F**). Thus, although increased NS selectivity observed at the individual neuron level is likely to aid in natural scene discriminability, it appears NS selectivity alone is not sufficient to improve natural scene discriminability. A strong shift in the NG preference index also appears to be required. To confirm this relationship, we compared the median NG preference index of all animals across the three rearing conditions with decoding accuracy, on an animal by animal basis. The correlation was significant (**Fig. 4G**), thus establishing that NG preference is predictive of natural scene discrimination.

Given the observation that the majority of neurons strongly driven by natural scenes were weakly responsive to gratings in standard-reared animals, it is expected that NS-only responsive neurons as well as neurons with a high NG preference index would outperform neurons with a low NG preference index in natural scene decoding. We verified this expectation by comparing natural scene decoding accuracy of neurons with NG preference indices greater than 0.5 to neurons with NG preference indices below 0. On an animal by animal basis, responsive neurons were pooled into two preference categories and accuracy was compared (**Fig. 5A**). As anticipated, we found decoding accuracy was lower for neurons with NG preference indices lower than 0. Analysis of the confusion matrices (**Fig. 5B**) revealed that similar to the comparison between standard-reared and dark-reared animals (**Fig. 3C,D**), the probability of error for both cross scene-type and within scene-type was lower in standard-reared animals compared to dark-reared animals. Furthermore, qualitatively the difference in performance was greatest for cross scene-type pairwise comparisons, as was the case in the standard-reared versus dark-reared comparison **(Fig. 5C,D)**. These results indicate that neurons with high NG preference index values contain information useful for discriminating the more similar scene pairs as well as the most different scene pairs.

**Figure 5.**
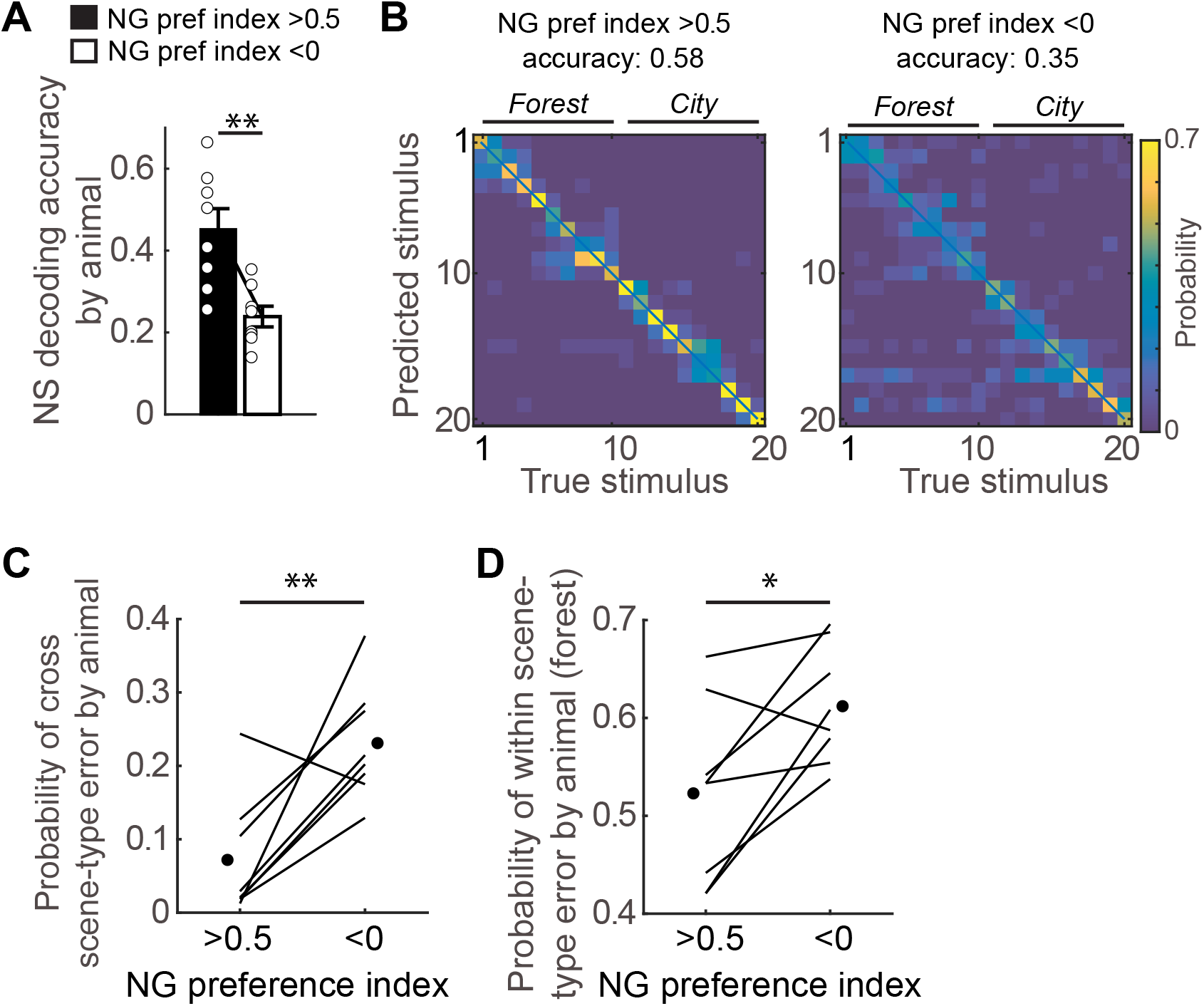
Neurons well-driven by grating stimuli performed poorly on natural scene discrimination. **A** In SR animals (n= 8 animals), grating-preferring neurons (NG preference index <0) were selected for decoding natural scenes and compared to natural scene-preferring neurons (NG preference index > 0.5). Decoding accuracy was significantly lower in the grating-preferring group (paired t-test, p=0.002). The number of neurons was matched across the two groups on an animal-by-animal basis (neuron number by animal: 127,154, 106, 181, 139, 147, 108, 118). **B** Example confusion matrix plots from one animal, 181 neurons. Accuracy is calculated as the sum of the diagonal (blue line). **C** Probability of cross-scene error (city and forest scene type values were averaged) was significantly higher in the grating-preferring group compared to natural scene-preferring group (Wilcoxon rank test, p= 0.007). **D** Probability of within-scene error, specifically for the forest scene type, was significantly higher in in the grating-preferring group compared to natural scene-preferring group (Wilcoxon rank test, p= 0.017). Error bars indicate S.E.M. *p<0.05, **p<0.01

Although we restricted our analysis to trials that contained eye movements less than 10 degrees, it is still possible that eye movements within this 10-degree limit were greater in DR or DVE conditions. Increased eye movements in either DR or DVE conditions could contribute to decreased discrimination accuracy. Therefore, we examined median eye movements by animal as well as pooled across animals (**Fig. S2E-F**). We found that movements were largely similar across animals, regardless of rearing condition. When pooling across animals we noted that eye movements in the SR condition deviated more from the median eye position than either the DR or DVE conditions. In addition, we found that pupil size was similar across conditions (**Fig. S2D,G**). Thus, differences in eye movements or pupil size specifically during stimulus presentation did not contribute to lower discriminability in either the DR or DVE conditions. In summary, these data demonstrate that increased preference for natural scenes can account for the higher discriminability observed in animals with normal visual experience compared to deprived animals.

### Development of natural scene preference and discrimination

Arousal state and overt movements substantially modulate neural activity in V1 [36–42]. Whether arousal-induced changes in neural activity serve to maintain or enhance stimulus representation during self-motion is an open question [43,44], and the influence of arousal on neural dynamics likely depends on the behavioral task [45,46]. Given that the animal is free to sit or run in our awake head-fixed recording configuration, it is possible that systematic differences between SR, DR, and DVE conditions in overall arousal state could contribute to the differences that we observed in natural scene discriminability. To address whether differences in arousal state throughout the imaging session may have impacted encoding of natural scenes and potentially influenced the accuracy of discrimination, we examined the extent to which pupil size, an indication of arousal state [47], and total locomotion during the entire session differed across conditions. We found that pupil size was similar across the three conditions (**Fig. 6A**). However, we did note that session-wide locomotion was systematically higher in the DR and DVE conditions relative to the SR condition. The time spent running was greater in DR and DVE conditions compared to SR. On average SR mice ran for 15±0.03% of the time, while DR mice ran 21±0.03% of the time, and DVE mice ran 25±0.02% of the time (**Fig. 6B**). In addition, the average locomotion speed during running bouts was increased in DR and DVE conditions relative to SR (**Fig. S2H**). Therefore, we cannot rule out the possibility that a 6-10% increase in locomotion induced changes in neural activity that outlasted the running epochs and contributed to the lower decoding accuracy that we observed in DR and DVE conditions.

**Figure 6.**
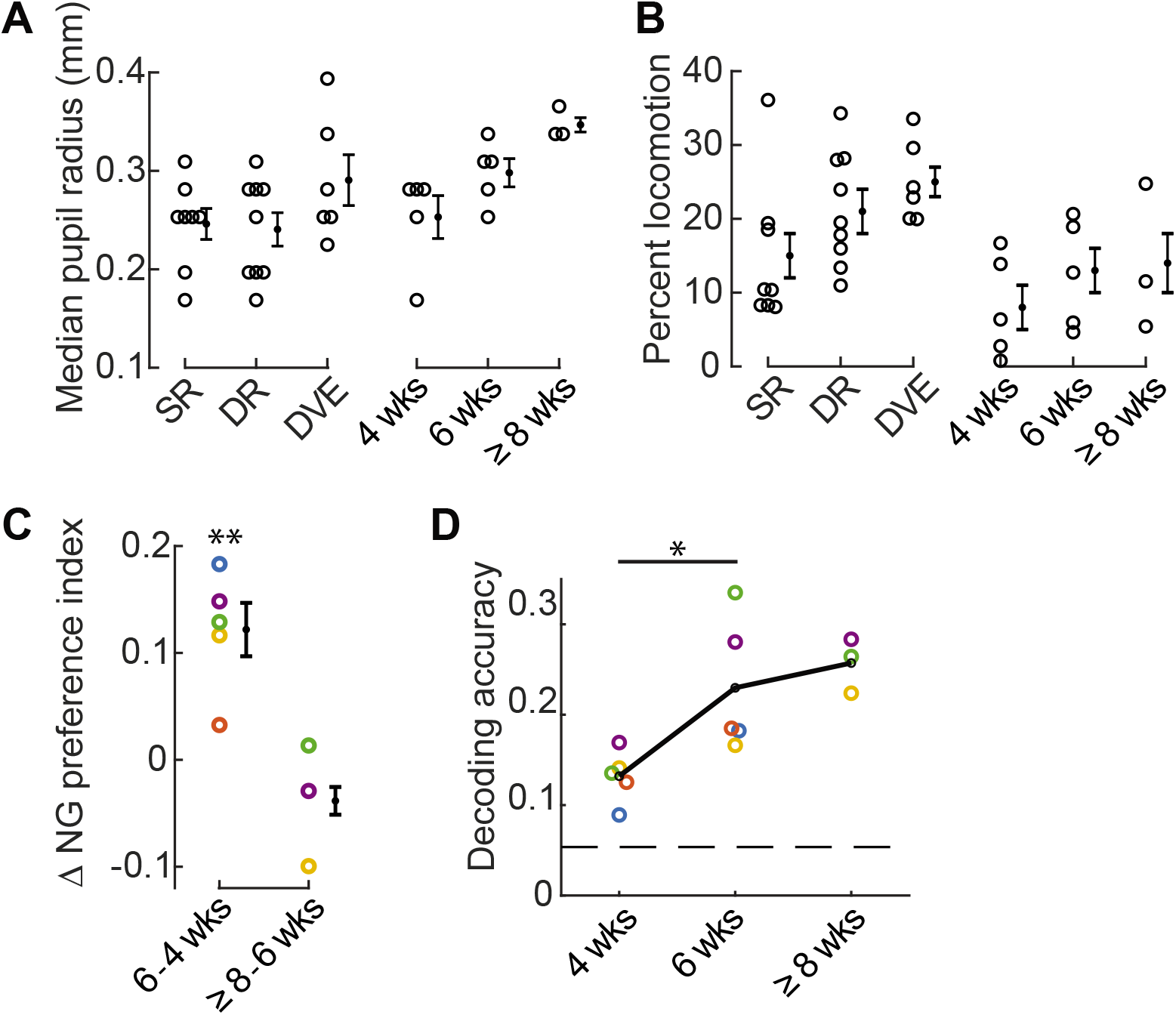
Development of natural scene preference and discrimination in relation to arousal state. **A** Pupil size of individual animals was largely overlapping across conditions, including SR, DR, DVE, and in the developmental SR time course spanning 4 to ≥8 weeks. **B** The amount of time individual animals spent running throughout the imaging session tended to be lower in the standard-reared condition. **C** The change in the median NG preference index for each animal is labeled by color. Mice color coded in blue, yellow, and orange were not treated with Doxycycline; mice colored coded by green and purple carried the SLC17a7cre allele. The difference between 4 and 6 weeks was significant, paired t-test p= 0.008. **D** Decoding accuracy of individual animals, labeled as in ‘C’, significantly increased between 4 and 6 weeks of age, paired t-test p= 0.029. Only animals with at least 16 trials for each natural scene stimulus were included.114 neurons per animal were used. Chance performance is 0.05 (dashed line). Error bars indicate S.E.M. across animals. *p<0.05, **p<0.01

Based on these results we sought to identify conditions in which we might observe low levels of locomotion coinciding with low discrimination accuracy, or a difference in discrimination accuracy that did not coincide with a change in locomotion. Five standard-reared mice were imaged at two different ages. The first time point was acquired at P28-29, the peak of the critical period for ocular dominance plasticity and within the critical period for binocular matching of ocular input [48]. The second time point was acquired two weeks later. We found that in standard-reared animals, the percent time spend running was comparable across these two ages, although was slightly higher in the older age (**Fig. 6B**). Pupil size during the imaging session was slightly larger at 6 weeks compared to 4 weeks of age (**Fig. 6A**). Next we computed the NG preference index and discrimination accuracy. We found that between 4 and 6 weeks of age the NG preference index significantly shifted towards higher values (**Fig. 6C**) and discriminability improved over this same time period by 75% (**Fig. 6D**). Taken together, these results demonstrate that when considering the 4 conditions studied, SR, DR, DVE, and SR development, the shift in NG preference index and improved discrimination accuracy can be de-coupled from session-wide differences in locomotion and pupil size.

We were able to successfully perform a third imaging session at 8 weeks of age for 3 of the 5 mice in the SR development condition (**Fig. 6C,D**). In contrast to the change between 4 and 6 weeks, we did not detect a large shift in the NG preference index, nor did we see a substantial improvement in natural scene discriminability. In summary, our time course analysis in standard-reared animals establishes that natural scene discriminability continues to improve after the peak of the critical period for ocular dominance plasticity.

### Response tuning and natural scene discrimination

Next we considered the impact of experience on neural responses to grating stimuli. First we addressed whether the decrease in grating responsiveness found in standard-reared relative to dark-reared animals negatively impacted the ability of the V1 population to decode grating stimuli using a nearest-neighbor decoder. We found that decoding accuracy was actually improved, despite the experience-dependent loss of neurons responding to low spatial frequency grating stimuli (**Fig. 7A**). The improvement in grating decoding was accompanied by an increase in orientation selectivity, detected on an animal-by-animal basis (**Fig. 7B**) as well as the pooled population across animals (**Fig. 7C**). This improvement in grating decoding did not occur in DVE animals (**Fig. S6A**). Although DVE animals displayed a decrease in the number of grating responsive neurons compared to dark-reared animals (**Fig. S6B,C**), an increase in orientation selectivity was not detected (**Fig. S6D)**.

**Figure 7.**
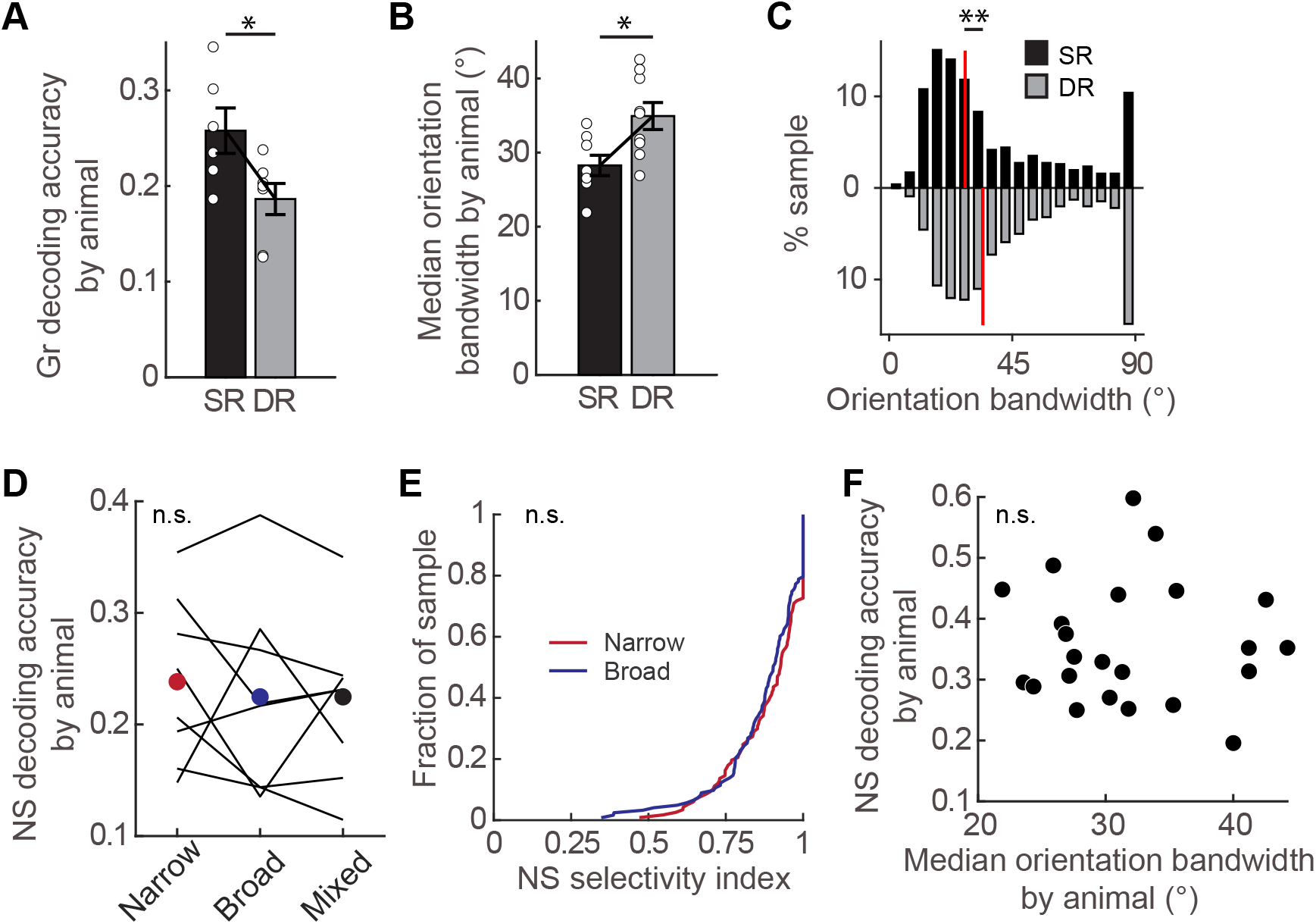
Experience-dependent refinement of orientation tuning was dissociated from natural scene discrimination. **A** Decoding accuracy of grating stimuli was significantly decreased in DR animals (n=7 animals) compared to SR animals (n= 6 animals; t-test p= 0.024). Only animals with at least 11 trials for each grating stimulus were included. 151 neurons per animal were used. Chance performance is 0.017. **B** Median orientation bandwidth across animals was significantly higher in DR animals (n= 9 animals) compared to SR animals (n= 8 animals; t-test, p= 0.011). **C** Orientation bandwidth values of individual neurons pooled across animals was significantly higher in dark-reared animals (1098 neurons from 9 animals) compared to standard-reared animals (770 neurons from 8 animals; KS-test, p= 5.17e-10). **D** Natural scene decoding accuracy of SR animals (n= 8 animals, lines) for each orientation tuned category as indicated. No difference between tuning categories was detected (repeated measures ANOVA, p= 0.755). Neurons with an orientation bandwidth less than 25° were considered to be narrowly tuned, and those more than 30° were considered to be broadly tuned. The number of neurons for a given animal was matched across tuning category, but varied by animal (neuron number by animal: 48, 34, 34, 42, 34, 30, 22, 28). **E** Natural scene selectivity index distribution of broadly (n=351 neurons) and narrowly (n=351 neurons) orientation-tuned neurons pooled across 8 SR animals that responded to at least one natural scene (KS-test, p= 0.081). **F** Natural scene decoding accuracy was not correlated with the median orientation bandwidth on an animal-by-animal basis. Pearson correlation coefficient r= −0.043, p= 0.847. Error bars indicate S.E.M. *p<0.05, **p<0.01

It has been proposed that experience regulates orientation tuning diversity at the population level such that natural scenes are effectively represented by a synergistic interaction between neurons broadly and narrowly tuned for orientation [11]. Therefore, we examined whether mixing broadly and narrowly tuned neurons from standard-reared animals improved discriminability relative to narrow or broadly tuned neurons alone. We found that mixing orientation selectivity did not improve natural scene discriminability compared to either narrowly tuned or broadly tuned neurons (**Fig. 7D**). Furthermore, neurons broadly tuned for orientation showed similar selectivity for natural scenes as did neurons narrowly tuned for orientation (**Fig. 7E**). These latter data directly demonstrate that orientation selectivity for grating stimuli was unrelated to the NS selectivity index. In contrast to the median NG preference index, the median orientation bandwidth did not correlate with natural scene decoding accuracy (**Fig. 7F**). This observation is consistent with our finding that narrowly tuned neurons did not out-perform broadly tuned neurons in natural scene decoding.

To further establish that neurons with a strong preference for natural scenes relative to grating stimuli function to enhance the discrimination and representation of natural scenes, we next examined the ability of individual neurons to discriminate natural scenes in relationship to each of the following five response properties: NG preference, NS selectivity, orientation selectivity, preferred spatial frequency, and spatial frequency selectivity. These are the five key properties that describe natural scene tuning as well as tuning to grating stimuli. A d-prime value for natural scene discrimination was computed for individual neurons. The preferred natural stimulus for each neuron was identified and the distribution of ΔF/F_0_ values in response to preferred stimulus trials was compared to the distribution of ΔF/F_0_ values to all other natural scene stimulus trials. Consistent with the decoding results above, the NG preference index was positively correlated with the natural scene d-prime (r=0.37 Spearman’s rank correlation p= <1e-50), and orientation bandwidth was not correlated with the natural scene d-prime (r=0.033 Spearman’s rank correlation p= 0.354; **Fig. S7**). Of the five response properties examined, only the NG preference index was positively correlated with the natural scene d-prime. Taken together, although diversity in orientation tuning may be beneficial to natural scene processing, particularly in a limited resource scenario such as a small number of neurons [11], our results demonstrate that visual experience early in life improves natural scene representation and discrimination by increasing the proportion of V1 neurons that are responsive to higher-order features.

## Discussion

To examine the impact of experience on natural scene discriminability we made use of 2-photon calcium imaging which permits the detection of all neurons within a field of view, including non-responsive neurons [4]. We found that experience increased the number of neurons preferring natural scene features relative to grating-preferring neurons, and that this increase can account for improvements in natural scene discriminability observed in standard-reared animals compared to animals raised without visual experience. In addition, we found that visual experience improves grating stimulus discriminability. The improvement was accompanied by a decrease in the number of grating-responsive neurons. Thus, experience reduces the number of neurons required to effectively encode grating stimuli.

Two recent studies reported that a fraction of mouse V1 neurons in standard-reared adults are preferentially driven by complex features found in natural images [21,22], a phenomenon that is observed in several species [19,23]. Moreover, in ferrets during early postnatal development, structured spontaneous activity emerges to specifically reflect natural scene statistics, indicating that V1 gradually adapts an internal model of the statistical structure of the natural visual environment [20]. However, it was not directly tested whether experience was required in the studies cited above. Here we show that the number of neurons exhibiting a response preference for natural scenes is significantly reduced in animals lacking visual experience. Our results provide evidence that experience drives individual V1 neurons to become directly sensitive to complex features found in natural scenes. Furthermore, we demonstrated that visual experience must occur early in postnatal development for improvements in natural scene discriminability to proceed.

There are several possible reasons that we did not find supporting evidence that diversity of orientation tuning improves natural scene discriminability in upper cortical layer 2/3. First, our decoding methods are an approximation of the theoretical discriminability limit, and therefore are not as precise as using Fisher information in conjunction with receptive field mapping, as was used in [11]. Second, it is well established that experience-dependent plasticity differs across layers [49], and within layer 2/3. Indeed, instructive influences driven by grating stimulus overexposure are more prominent in deeper layer 2/3 compared to upper layer 2/3 [4]. Thus it is possible that the impact of orientation tuning diversity on natural scene discriminability is dependent on cortical depth.

Contextual modulation located outside of the classic receptive field can account for variations in natural scene feature selectivity of individual neurons [12,13,17,23,27] and may contribute to the enhanced discriminability in standard-reared animals we report here. For example, it was previously shown that normal development of neuronal sensitivity for natural scene statistics in the receptive field surround is prevented by dark-rearing [17]. Surround-modulation is well positioned to mediate these effects for shapes that extend outside of the receptive field and could involve feedback from secondary visual cortex [29,30] as well as for higher-order statistics such as the phases of the Fourier spectrum [17,27]. The experiments in the present study do not distinguish between these possibilities. The extent to which natural-scene preferring neurons develop from neurons tuned to low level features such as oriented bars of varying spatial frequencies, is unclear. Given that the total number of responsive neurons was the same in animals raised with or without visual experience and the loss of low spatial frequency responsiveness was associated with a gain in natural scene sensitivity in standard-reared animals, it is possible that natural scene-preferring neurons directly emerge from the low-spatial frequency preferring population.

It is becoming increasingly apparent that the mouse visual system can respond to high spatial frequencies in moving stimuli [34], consistent with natural visual behavior [50]. The development of selectivity for direction of motion is highly dependent on visual experience, observed at the cortical level [51–54] as well as at upstream stages such as the retina [55]. Furthermore, the representation of direction of motion may be more unstable on a neuron-by-neuron basis within V1 compared to the representation of orientation [56,57]. The use of static stimuli allowed us to demonstrate that experience-mediated improvements in scene representation are observed for stimuli that do not contain motion.

The initial formation of cortical receptive field properties [58–61], such as orientation selectivity to grating stimuli [60], as well as retinal response properties [62] do not require visual experience. Innate properties are subsequently shaped by experience as well as by development itself. For example, parvalbumin inhibitory neurons broaden their orientation selectivity from 3 to 4 weeks of age and this broadening requires visual experience. During this same time frame orientation tuning becomes sharper in excitatory neurons, even in the absence of visual experience [63]. Classic stripe-rearing in which the animal is over-exposed to a limited repertoire of stimuli is a more direct method to examine how the neural activity that is evoked by a defined stimulus refines neural responses in the visual cortex [64]. Refinement involves a passive loss of responses to the non-experienced stimulus [4,65,66] as well as an instructive component that drives an overrepresentation of the experienced stimulus [4]. These studies raise the possibility that experience-dependent refinement of low-level feature representation underlies improved natural scene representation in standard rearing conditions as well. This could in theory arise through refinement of orientation selectivity, preference, or increased responsiveness to high spatial frequency gratings experienced during natural vision [9,10]. Although we did detect an experience-dependent sharpening of orientation tuning, these changes were not correlated with improved natural scene discriminability.

As noted above, previous studies in the mouse did not detect experience-dependent sharpening of orientation tuning [63], see also [48]. Given that the present study was restricted to upper layer 2/3, cortical depth of recording, as well as differences in age and a larger sample size, may have contributed to our ability to detect an effect of experience on orientation selectivity. Notably, the proportion of V1 neurons responding to high spatial frequencies was not different between standard- and dark-reared conditions. As such it is unlikely that increased sensitivity to high spatial frequencies was responsible for the higher grating discriminability or the higher natural scene discriminability that we observed in upper layer 2/3 of standard-reared animals.

In summary, our results demonstrate that natural scene discriminability continues to develop past the peak of the critical period for ocular dominance plasticity, requires early visual experience, and might not rely on the refinement of low-level feature representations.

## Acknowledgements

We thank Alison Barth and Doug Ruff for useful discussions, Tom Prigg, Jake Brezinsky, Jeff Good, Alex Hsu, and Jorge Reyes-Arbujas for technical assistance.

## Disclaimers

The views and conclusions contained herein are those of the authors and should not be interpreted as necessarily representing the official policies or endorsements, either expressed or implied, of IARPA, DoI/IBC, or the U.S. Government (TSL and SJK). The U.S. Government is authorized to reproduce and distribute reprints for Governmental purposes notwithstanding any copyright annotation thereon.

## Author Contributions

NK, PS, BBJ, and TF collected the data; NK, JK, PS, BBJ, TF, SMC, and SJK analyzed the data; NK, JK, PS, BBJ, SMC, SJK, and TSL designed the experiments and wrote the manuscript.

## Methods

All experimental procedures were compliant with the guidelines established by the Institutional Animal Care and Use Committee of Carnegie Mellon University and the National Institutes of Health. For imaging experiments, to express the green fluorescent protein calcium indicator GCaMP6f selectively in excitatory neurons [67], homozygous EMX1cre mice (Jackson Laboratories, stock no 005628) were crossed with homozygous Ai93-heterozygous Camk2a-tTA mice (Jackson Laboratories, stock no 024108), except for two animals (**Table S1**), which were offspring from a cross of homozygous SLC17a7cre mice (Jackson Laboratories, stock no 023527) crossed with homozygous Ai93-heterozygous Camk2a-tTA mice. All experimental mice were heterozygous for all three alleles. Mice were housed in groups of two either under a 12 /12 hour reversed light cycle or in a ventilated dark cabinet, with the exception that three experimental mice were single housed post-surgery due to an uneven number of mice in the initial cage. Dark-reared mice were placed in the light-tight cabinet at age P6. During dark-rearing, daily health checks were performed under red led (less than 2 lux); exposure to the red light was limited to 5 minutes or less. Mice had ad libitum access to food and water unless noted. EMX1cre-containing mice were treated with doxycycline as in [56] to prevent aberrant activity [68], except in two DR animals and 3 SR animals for which imaged started at 4 weeks of age (**Table S1**). The mice not treated with doxycycline used in this study did not exhibit signs of aberrant activity (**Figure S1H-J**).

### Animal preparation

During surgery, mice were anesthetized with isoflurane (3% induction, 1-2% maintenance). Core body temperature was monitored and maintained at 37□°C using a feedback heating system. A stainless-steel bar, used to immobilize the head during imaging, was glued to the right side of the skull and secured with dental cement. A 3-mm-diameter craniotomy was made over the primary visual cortex in the left hemisphere, identified by coordinates and landmarks as described in [69,70]. The craniotomy was then covered with a double glass assembly and sealed with dental cement.

### Go/No-go behavioral task

A 3D-printed custom acrylic lickport was used to record licks and to deliver water rewards to mice that were water restricted to 750 μl/day. Licks were recorded by the tongue breaking an infrared light path between LED optical switch photodiodes (Vishay Semiconductors; TCZT8020-PAER). Water rewards were delivered by gravity using a 3-port solenoid valve that opened for a defined period of time following a beam-break on ‘Go’ trials (Lee Company; LHDA1231115H) through a 0.02-inch-diameter stainless steel tube. In order to prevent the pooling of water on the lickport, excess or pooled water was pumped away from the steel reward tube by plastic tubing using a peristaltic pump (Fisherbrand; 70730-064). An mBed microcontroller processor (mBed 1768 Demo Board; Mouser 7711-OM11043598) and custom scripts were used to schedule ‘Go’ and ‘No-go’ stimuli, sample lick times at a frequency of 10 Hz, and gate the release of water rewards. The volume of the water delivered per beam beak was controlled by the duration that the solenoid valve was open, the rewarded volume was ~6-7μL. During behavior, mice were mounted on stainless steel bar positioned over a Styrofoam ball floated on a cushion of air. Visual stimuli were presented on a screen (Dell; 30”, 2560×1600 resolution; 9TDTX) that was positioned 25 cm away from the mouse in front of the right eye, angled at 50° with respect to the midline of the animal.

Stimulus trials were separated by an inter-trial-interval of 2 seconds during which a black screen was presented. For every ‘Go’ trial that the mouse licked, one water drop was delivered. False alarms (licking on a ‘No-go’ stimulus) were punished with a timeout: the black screen inter-trial-interval duration was extended for 1 to 8 seconds, depending on the mouse (some responded well to 1 second, while others required a longer duration). If the mouse licked at any point during the timeout, the timeout duration was reset and triggered again. Misses (a failure to lick on the ‘Go’ stimulus) were not punished. Mice performed one session per day. Go stimulus trial frequency within a session was set to 35%. Performance was calculated as follows:

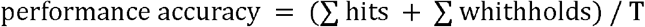

where *T* is the total number of trials, a hit is at least one lick on a Go stimulus trial, and a withhold is when the mouse did not lick on a No-go stimulus trial. Hit rate and withhold rate were calculated as follows:

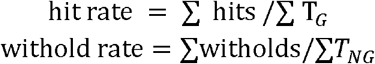

where *T_G_* is a Go stimulus trial, *T_NG_* is a No-go stimulus trial. A total of 5 static ‘City’ natural scene images were used. Natural scene stimulus ID #16 (**Fig. S1B**) was set as the No-go stimulus for the duration of the experiment. During shaping, an image that was 25 frames distal to stimulus ID#16 was selected from the city movie sequence as the Go stimulus. Once animals achieved a performance of 85% or higher for 2 consecutive days during the initial shaping, the Go stimulus was updated to be more similar to stimulus ID#16 and the new Go stimulus tested for 3 sessions. The best performance out of the three test sessions for a given image was reported in Figure 1C.

### Stimulus presentation during imaging

A total of 80 stimuli were presented as static images in Matlab (Mathworks, Boston, MA) using Psychophysics Toolbox (http://psychtoolbox.org) on a screen positioned 25 cm away from the right eye (contralateral to the imaged hemisphere) angled at 50° with respect to the midline of the animal. The size of the screen was 64cm x 40cm, subtending 146° x 92° of visual angle. Two categories of natural scenes that elicited strong neuronal responses were used: scenes from a “Forest” video and scenes from a “City” video downloaded from: www.youtube.com/watch?v=uEb_0Q1uR3o, www.youtube.com/watch?v=qIWSbBXmaj8. Ten frames from each movie were selected and converted to grayscale, matched for luminance, and displayed at full screen. SSIM was computed using the ‘ssim’ function in Matlab. Power spectra of natural scenes in Figure S1C were computed by extracting 100 400 x 400 pixel patches (17° x 17°) were from each image and the power spectrum of each patch computed, with 10 cycles per degree as the maximum frequency. The power along eight radials (22.5° spacing) was averaged and plotted separately for City and Forest scenes for spatial frequencies up to 1 cycle/°. To compute the difference image shown in Figure S1C inset, 100 400 x 400 pixel patches (17° x 17°) were extracted for each image and their power spectrum computed, with 10 cycles per degree as the maximum frequency. The dc-center 40 x 40 patch (corresponding to 1 cycle per degree maximum frequency) was then extracted from each power spectrum and averaged across all the patches from city scenes and for forest scenes, respectively.

Sinusoidal gratings were shown at an aperture of 60° on a gray background, 100% contrast, 12 orientations (15° spacing), 5 spatial frequencies (0.05, 0.1, 0.2, 0.3, 0.4 cycles/°). The aperture edge was smoothed with a Gaussian blur (α=10 pixels, corresponding to 1°) to eliminate sharp edges. Grating and natural scene stimulus sets were presented in alternating blocks of approximately 10 ‘runs’ for the natural scene set and 5 ‘runs’ for the grating set, where a ‘run’ included one presentation of each stimuli for a given set. All stimuli were randomized and interleaved with a gray screen. Stimulus duration was 1 second, and the gray screen duration was 2 seconds. We verified that our natural scene stimulus set contained a range of orientation concentration indices (OCI; **Fig. S1E-G**). OCI values were derived from 20° patches, calculated as in [11].

### Data acquisition

Mice were head-fixed atop a freely rotating air lifted Styrofoam ball. Prior to imaging, mice were given 15 minutes to acclimate to the movement of the ball. Calcium imaging was performed using a resonant, two-photon microscope (Neurolabware, Los Angeles, CA) controlled by the Scanbox acquisition software (Scanbox, Los Angeles, CA), and collected at a depth of 150-250 microns below the pia. The light source was a Coherent Chameleon Ultra II laser (Coherent Inc, Santa Clara, CA) running at 920 nm; green emissions were filtered (Semrock 510/84-50) and captured with a PMT (Hamamatsu H1 0770B-40) and amplifier (Edmund Optics 59-179). The objective was a 16x water immersion lens (Nikon, 0.8 numerical aperture). Data were collected from a 440um X 550um field of view at a frame rate of 15.48 Hz. Locomotion and pupil movements of the left eye were recorded (Dalsa Genie M1280) and synchronized with the calcium imaging frames. To compute locomotion, a threshold on the luminance intensity of the treadmill motion images was applied and the phase correlation was computed between consecutive frames to estimate the translation between the frames. The estimated translation was converted to movement speed, taking into account the dimensions of the treadmill motion images (7.8□cm□×□7.8□cm) and the acquisition rate of the camera (30□Hz). To align ball motion data with calcium imaging data, the raw movement speed was down-sampled to match the acquisition rate of calcium imaging. Stimulus epochs that included a treadmill speed of greater than 2 cm/ second for more than one frame were considered to contain locomotion and removed from further analysis Pupil location was estimated from eye-tracking videos using a circular Hough Transform algorithm (**Fig. S3B**).

### Pre-processing of calcium signal

Raw data image sequences were pre-processed with Suite2p toolbox. Pre-processing included frame alignment to correct for motion, segmentation, and neuropil correction [71]. In Suite2p, the latter two steps are achieved iteratively using a model where the recorded signal consists of activity of the ROIs, the neuropil and measurement noise. The neuropil signal is represented in a set of spatially-localized isotropic 2-dimensional raised cosine basis functions, which allow the signal to vary slowly across space. After model estimation, neuropil contamination was removed by subtracting a scaled-down version of the neuropil signal using estimated scaling factors (the estimation was typically between 0.5-1.0). All detected segments were further curated by a classifier trained with data collected in-house. After these automated steps, labeled segments were confirmed, rejected, or selected by manual visual inspection; less than 5% of the segments were rejected and/or selected.

For each segmented neuron, the trial-averaged response to a given stimulus was computed by averaging the ΔF/F_0_ across the entire 1 second presentation of that stimulus. ΔF/F_0_ was calculated as:

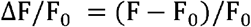

where F is the Suite2P-generated neuropil-corrected fluorescent signal of a given neuron for a single frame in the series, and F_0_ is the global baseline, calculated as the mean fluorescent signal across all gray screen presentations in a given ‘run’ excluding the top 25^th^ percentile of values.

Sampled time points (calcium imaging frames) that contained large cross-frame alignment translations (greater than 15 microns) or a drop in signal intensity (greater than 3 standard deviations) is an indication of z-motion and were marked as non-informative and removed from the time series. In addition, sampled time points associated with eye movement > 10° from the median position, or locomotion > 10 cm/s were also removed. Calcium ΔF/F_0_ signals were interpolated over removed frames in the case that the number of consecutive non-informative frames was 2 or less for a given stimulus trial. Trials were excluded from further analysis if > 2 consecutive frames were removed, or if more than 10% of the frames were removed.

### Data analysis

To quantify the total number of neurons imaged in a session, including non-responsive neurons, we chose to use an objective method. The goal of this inclusion criterion was to ensure that all segments exhibited activity. The calcium signal of each segment generated using Suite-2P was deconvolved [56], and segments with at least one inferred event count larger than 6 events within a 200 ms time bin were considered to be neurons. For each segment *n*, spike events e_*n*_ were inferred from fluorescence f_*n*_ using the following model:

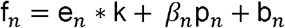

where k is the temporal kernel and b*_n_* is the baseline fluorescence. A single kernel was derived for all neurons in one imaging session, all other parameters were unique to each individual neuron. Neuropil, which is a contamination of the fluorescence signal f*_n_* from out of focus cell bodies and nearby axons and dendrites, is modeled by two parameters: p*_n_*, the time course of the neuropil contamination, and β*_n_*, the scaling coefficient. * denotes convolution. Using this model, e*_n_*, k, *β_n_*, and b*_n_* were estimated by a matching pursuit algorithm with L0 constraint, in which events were iteratively added and refined until the threshold determined by the variance of the signal was met.

To determine whether a neuron responded significantly to a given stimulus, a permutation test was used (5000 iterations). A null-distribution was generated using the last second of the inter-leaved gray screens and the median response to a given stimulus compared against the null-distribution. Estimated p-values were adjusted for multiple comparisons using false discovery rate (FDR). Alpha was set to 0.01. The number of neurons in each session that passed this criterion is indicated in Table 1, column 4. Neurons with a significant response to at least one of the 80 stimuli presented (60 grating and 20 natural scene stimuli) were considered visually responsive.

### Grating tuning

Orientation and spatial frequency preference were determined using a two-dimensional Gaussian model, fit to single trial responses. Only neurons responding to at least one grating stimulus were included. A nonlinear least-squared regression model was fit for each neuron such that the observed single trial response, R, was a function of the orientation θ and the spatial frequency φ of the presented stimulus of the form:

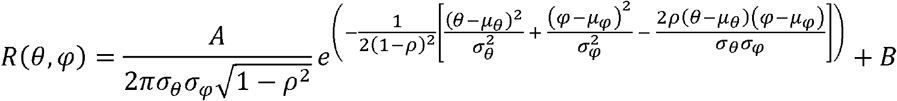

where μ_θ_ is the estimated preferred orientation and μ_φ_ is the estimated preferred spatial frequency of the neuron, and σ_θ_ and σ_φ_ describe the widths of tuning to those parameters. Prior to fitting, the preferred orientation was estimated by averaging the response, *F* across all spatial frequencies for a given stimulus orientation, θ and calculating half the complex phase of the value

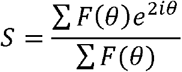

[33,63]. The preferred spatial frequency was estimated by selecting the spatial frequency that generated the maximal significant response at the estimated preferred orientation. The covariance of responses for orientation and spatial frequency is captured by the correlation term ρ. The parameter A is the amplitude and B is the baseline response of the cell. For fitting, the lower and the upper bound of allowed values for μ_φ_ was set by the range of the presented stimuli, which was 0.05 to 0.40 cycles per degree. The lower bound for σ_θ_ and σ_φ_ was set at to be positive to prevent fits with zero or negative widths (0.001 and 1.0 cycles/°, respectively). The bandwidths of the Gaussian tuning were described using half-width at half-maximum (HWHM). The HWHM bandwidths for both orientation and spatial frequency were calculated as:

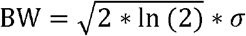

where *σ* is the width parameter of the Gaussian fit. The preferred orientation was estimated by calculating the complex phase, and the preferred spatial frequency was estimated by selecting the spatial frequency that generated the maximal significant response. To select neurons well-tuned to grating stimuli, the chance distribution of R^2^ was calculated from fitting the above model with permuted stimulus labels on individual trials, 1000 times for each neuron. Neurons whose R^2^ exceeded the 95^th^ percentile of the chance R^2^ distribution were accepted as tuned to grating stimuli. The median orientation bandwidth of tuned neurons in standard-reared animals was 27.8°. Based on this observation, neurons with an orientation bandwidth less than 25° were considered to be narrowly tuned, and those more than 30° were considered to be broadly tuned.

### Natural scene tuning

The NG preference index was calculated as:

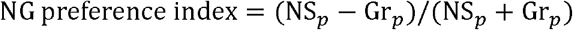

where NS_p_ is the preferred natural scene trial-averaged signal (one out of 20 possible stimuli) and Gr_p_ is the preferred grating trial-averaged signal (one out of 60 possible stimuli).

To compute the NS selectivity index, we first normalized the trial-averaged responses (*R*) for each natural scene stimulus (*i*) that elicited a significant response for a given neuron as:

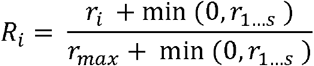

where *s* is the number of stimuli presented (20 natural scenes in our case), and *r* is the trial-averaged mean. We then computed the NS selectivity index as:

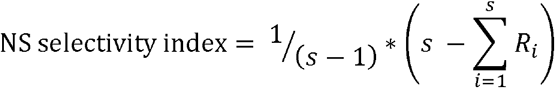

The intuition is that the smallest area under a normalized tuning curve is proportional to the stimulus selectivity of the neuron. If the neuron responds to one stimulus only, the NS selectivity index value will be 1, and if the neuron responds equally to all stimuli, the NS selectivity index value will be 0. We computed the standard error of the mean (SEM) using a bootstrap method, where n was the number of repetitions of each stimulus. For each stimulus, the neuron’s response to that stimulus was sampled n times with replacement and the NS selectivity index was computed from the re-sampled trials. This was repeated 1000 times to generate a distribution of NS selectivity index values. The standard deviation of this distribution is the SEM. Only neurons that responded to at least one natural scene and had a NS selectivity SEM < 0.1 were included.

### Population decoding

We used 2 distinct classifiers to decode the presented stimuli. First, k-nearest neighbor classifiers were used to decode the visual stimulus identity from the vector of single trial responses, by using the distance between response vectors. In our case, the k-nearest-neighbor classifier estimated the stimulus identity for a given response vector by identifying the most frequent stimulus identity of its k closest response vectors. To identify the nearest neighbors for a given response vector, *r_t_*, we used correlation distance to other response vectors, *r_n_* [22]:

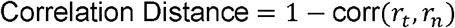

For each session, data was divided so that a single set of response vectors consisted of one trial of each stimulus. This resulted in the number of sets being equal to the number of trials that each stimulus was shown. During decoding, the possible neighbors for a test response vector consisted of all response vectors not belonging to the test set. This ensures an unbiased representation of possible nearest neighbors across stimuli. This process was repeated across each response vector and each set. We reported the performance of this decoding process as accuracy across all response vector tested. Only the neurons that were responsive to grating stimuli were included in decoding. To compare accuracies across animals, we matched the number of neurons used in classification to the minimum number of neurons. To determine the optimal value of k, the number of neighbors, we performed decoding on data from 7 animals collected in a separate pilot experiment, sweeping k from 3 to 15. The data from the pilot experiment were not used in any other part of this study, ensuring a properly cross-validated estimate of k. We ranked the values of k that yielded the highest accuracy for each animal. We found k= 3 had the best average rank across animals. Therefore, we used k= 3 to decode grating and natural scene stimuli.

Second, a Naive Bayes classifier which is a probabilistic classifier that finds the maximum a posteriori estimation using Bayes’ Theorem and the assumption of conditional independence was used:

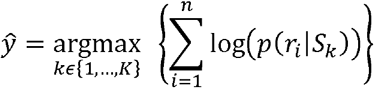

where 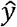 is the class label, *K* is the total number of stimuli, *n* is the total number of neurons, and *r_i_* is the response. We ignore the prior probability of each stimulus, as we down-sampled trials in the analysis such that each stimulus was observed an equal number of times. As is typical in calcium imaging recordings, fluorescence responses were asymmetric and sparse. To more accurately capture the true response distribution, for each neuron we quantized the marginal response of the neuron into N equally probable bins, and estimated the probability of response for each bin and stimulus. The number of bins, N, was optimized for natural scene decoding performance using the same pilot data set described above; N was determined to be 3. We confirmed that the optimal bin number (median across animals) used in this study was 3; this was the case for both natural scene and grating decoding. We report the performance on the heldout data from leave-one out cross validation.

In both the k-nearest neighbor and Naïve Bayes decoders, trial number was set to be the minimum trial number across all stimuli and animals, and the number of neurons used was set to be the minimum across animals. Chance level of both decoders was 1/N_s_, where N_s_ is the number of stimuli, i.e., N_s_ was 60 for the grating stimulus set an N_s_ was 20 for the natural scene stimulus set. The above three parameters are reported in the figure legends.

### Individual neuron discriminability

To quantify the discriminability of individual neurons, d-prime was computed as:

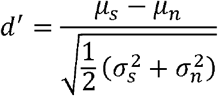

where the signal and noise distributions with mean and standard deviations were represented as μ_s_ and σ_s_, and μ_s_ and σ_s_, respectively. The signal was defined as the neuron’s response across trials to the preferred natural scene stimulus, and the noise was defined as the neuron’s response across trials to all other natural scene stimuli (non-preferred). The preferred natural scene stimulus was defined as the stimulus that elicited the maximal trial-averaged signal. trial number was set to be the minimum trial number across all stimuli and animals, 24 trials in this case.

### Statistical analysis

Statistical tests were performed using SPSS (IBM), expect for permutations and FDR, in these cases Matlab was used. Error is reported as S.E.M., defined as the standard deviation divided by the square root of the sample size. Non-parametric tests were used in cases data were not normally distributed; normality was assessed using the Shapiro-Wilk test. Alpha was set to 0.05 unless noted in Methods Details. Information regarding the statistical test used, p value, the exact value of the sample size n, and what n represents (animals or neurons) can be found in the legends. One animal in the DR condition was excluded because eye tracking failed during data collection. Litters were randomly assigned to experimental conditions.

