## Supplemental Information for "Development of natural scene representation in primary visual cortex requires early postnatal experience"

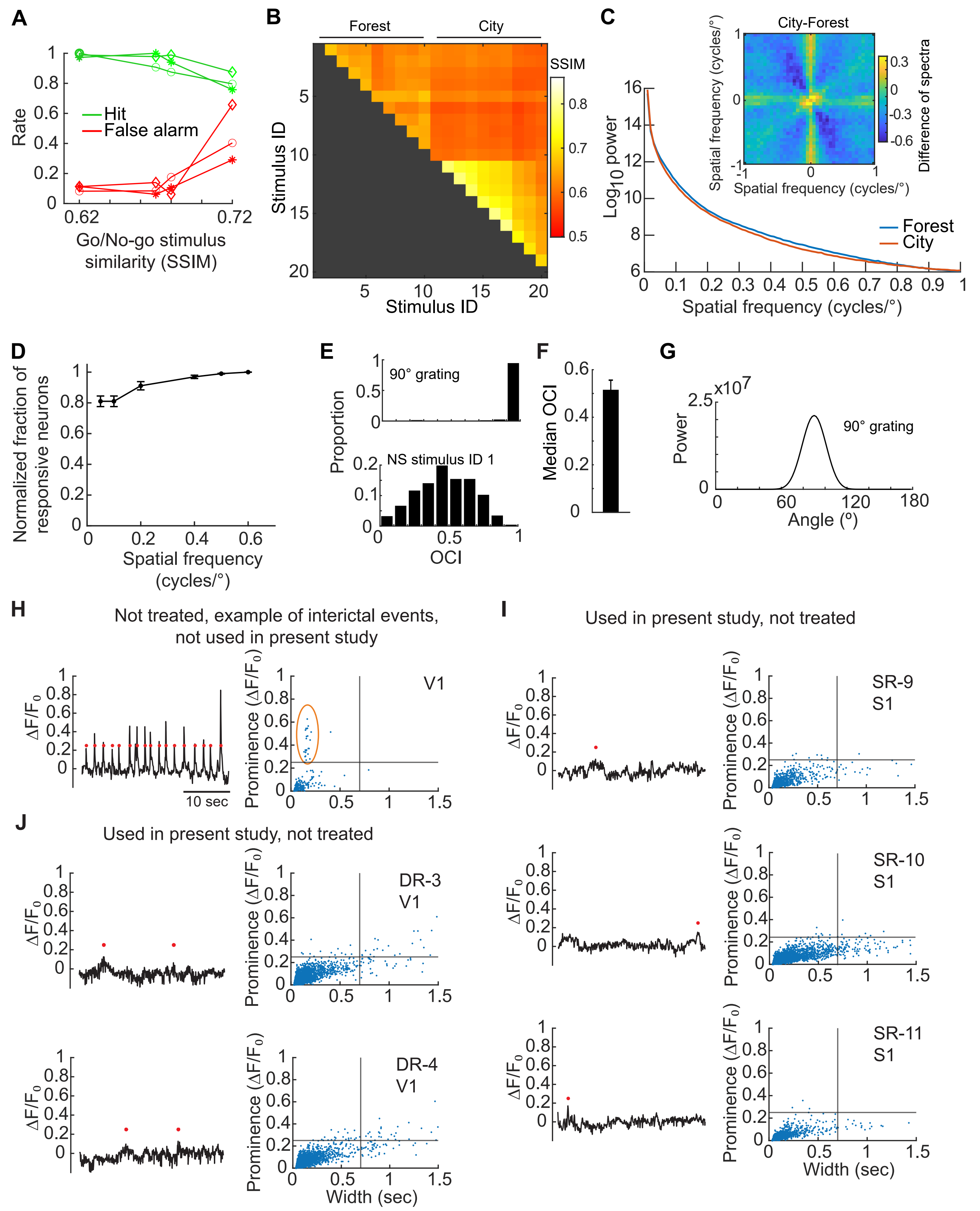

**Figure S1. Characterization of stimulus sets used in this study, including a quantitative description of natural scenes, related to Figure 1 and Methods.**

**A** Go/No-go behavioral task accuracy expressed as hit (correct licks) and false alarm (incorrect licks) rate.

**B** Pairwise structural similarity (SSIM) is plotted for all scene pairs. Individual frames from two different movie sequences were selected such that frames close together in the sequence were more similar. The relative frame spacing between the first frame and the subsequent frames for both the Forest scenes and City scenes are as follows 0,2,4,6,8,9,10,12,14,16. (cont.)

**C** The mean power spectrum of Forest scenes and City scenes, averaged across 8 orientations. The spectral power of natural scene stimuli was strongest for lower spatial frequencies, regardless of scene type. Inset, the difference between the power spectrum of city and forest scenes shows that city scenes had more power in the cardinal orientations relative to forest scenes and were weaker in the off-cardinal orientations. This is because city scenes have more vertical and horizontal structures, and less rounded structures relative to forest scenes.
**D** Saturation curve in which the mean fraction of responsive neurons across animals (n= 4 standard-reared animals) is plotted relative to stimulus spatial frequency. Fraction of neurons was normalized to the cumulative number of responsive neurons per animal.
**E** Top, example distribution of orientation concentration indices (OCI) derived from 20° patches sampled from a 90° orientated grating (0.05 cycles/°). Bottom, distribution of OCI values derived from stimulus #1 of the natural scene stimulus set.
**F** Median OCI across all 20 natural scenes.
**G** Mean filter power of the same 90° grating as in 'E'.
**H** Example of interictal events observed in an animal not included in the present study. Left, time series of fluorescence, averaged across the full field of view. Peaks above 0.25 $\Delta F/F_0$ are indicated (red dots). Right, peaks having a similar width above 0.25 were detected (gold outline) and considered to be aberrant interictal events as defined by [S1]. Images were acquired from primary visual cortex (V1).
**I** Two dark-reared animals used in this study were not treated with doxycycline, DR-3 and DR-4. Neither animal exhibited interictal events. Images were acquired anterior to V1, and included primary somatosensory cortex (S1). Labels as in 'H'.
**J** Three standard-reared animals (SR-9, SR-10, SR-11) included in the 4-8 week time course were not treated with doxycycline. Labels as in 'H'.
None of the animals in 'I' or 'J' exhibited interictal events. Error bars indicate S.E.M.

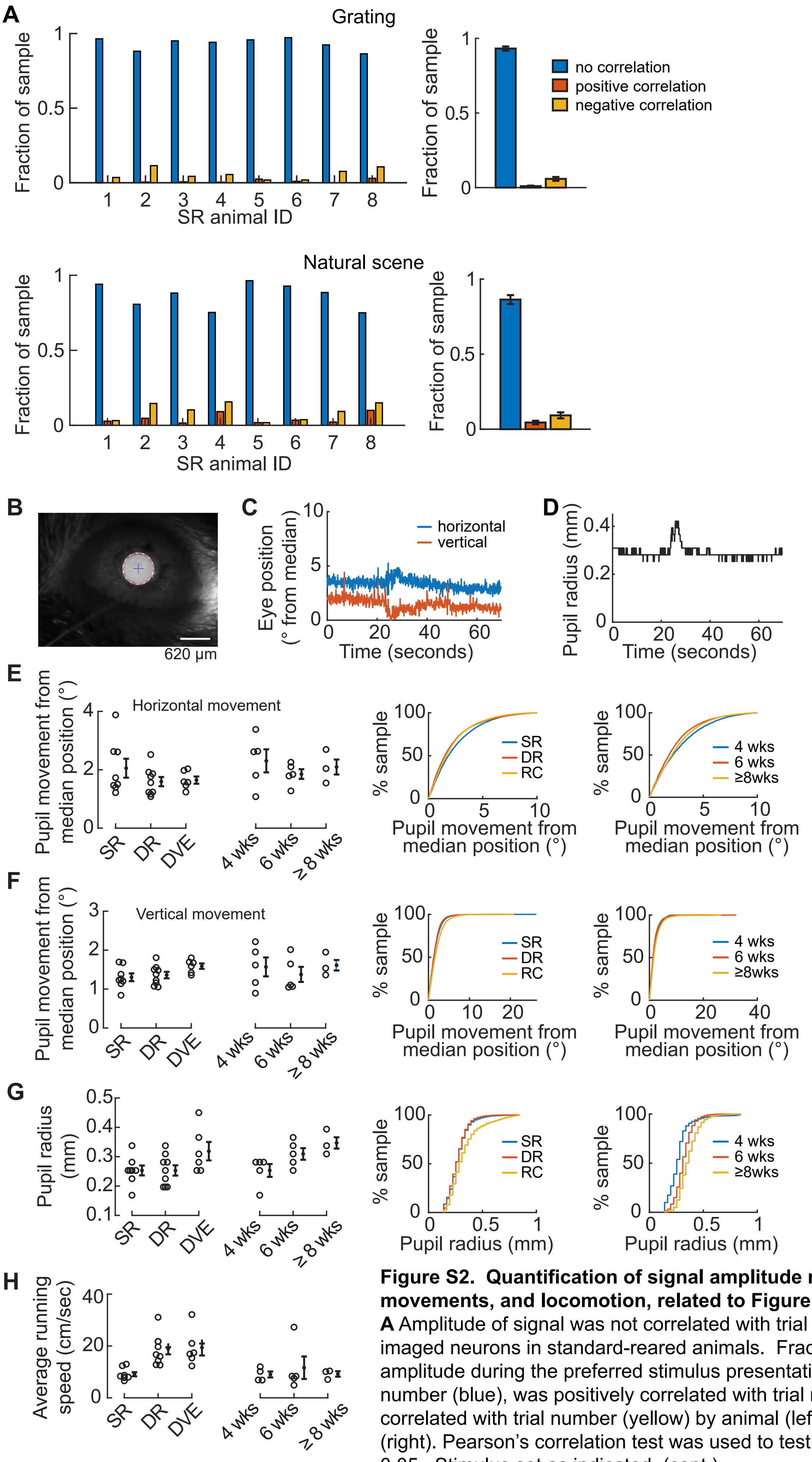


**Figure S2. Quantification of signal amplitude realtive to trial number, eye movements, and locomotion, related to Figure 1 and Methods.**

**A** Amplitude of signal was not correlated with trial number for the majority of imaged neurons in standard-reared animals. Fraction of neurons for which signal amplitude during the preferred stimulus presentation was not correlated with trial number (blue), was positively correlated with trial number (orange), and negatively correlated with trial number (yellow) by animal (left), and averaged across animals (right). Pearson's correlation test was used to test for significance, alpha was set to 0.05. Stimulus set as indicated. (cont.)

**B** Example eye-tracking frame depicting pupil centroid (+) and circumference (red-dashed circle) identified using the Hough transform algorithm.
**C-D** Pupil size and position tracked over time.
**E-F** Pupil coordinates specifically during stimulus presentation for those trials that meet the inclusion criteria described in the Methods such as no running, and no eye blinking.  Average movement by animal (right), and pooled across all tracking frames by experimental condition as indicated (center and right).
**G** Pupil size specifically during stimulus presentation, for those trials that met the inclusion criteria described in the Methods such as no running, and no eye blinking.  Average pupil size by animal (right), pooled across all frames, by experimental condition as indicated (center and right).
**H** Average running speed by animal, including both stimulus and gray-screen epochs and all trials.  This analysis describes the behavioral state during the full imaging session.
Error bars indicate S.E.M.

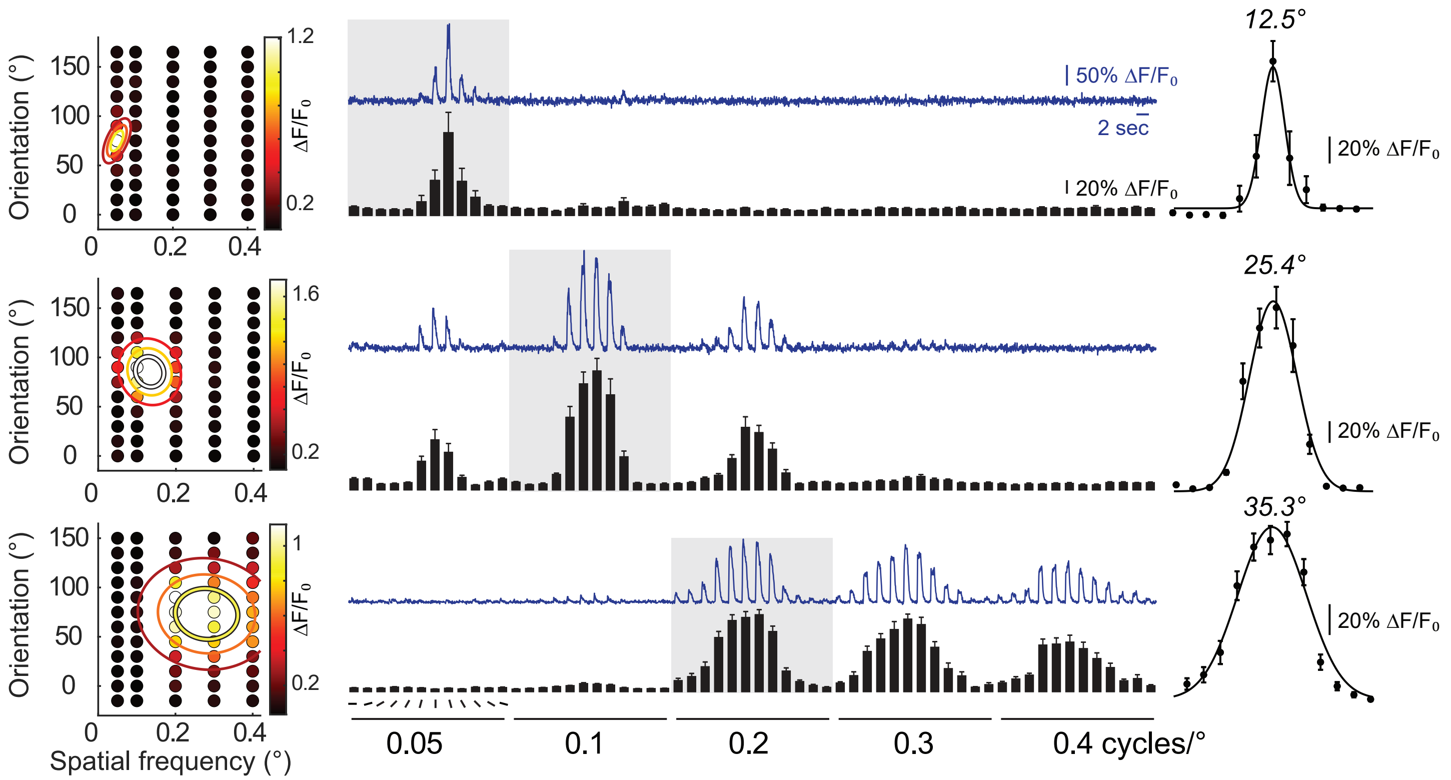


**Figure S3. Quantification of 2-dimensional tuning curves, related to Figure 2, 4, and Methods.**
Example 2- dimensional tuning curves, three neurons. Left column, two-dimensional Gaussian fits of the trial-averaged response. The grid of small circles represents the mean response to a given stimulus orientation and spatial frequency as indicated by the y-axis and x-axis, respectively. The three outlined ellipses of decreasing radius correspond to 75%, 50%, 25% of the peak Gaussian fit, respectively. Right column, blue traces are the average of 11-18 trials, traces were clipped to include 1 second of the visual presentation and 1 second of the following gray screen. Visual stimuli were randomly presented and re-sorted in this plot, as indicated by the lines in the lower left (orientation) and horizontal lines (spatial frequency). Black vertical bars indicate the mean response across trials. Gray shading indicates the spatial frequency and orientation angle of stimuli plotted in the tuning curve on the far right.

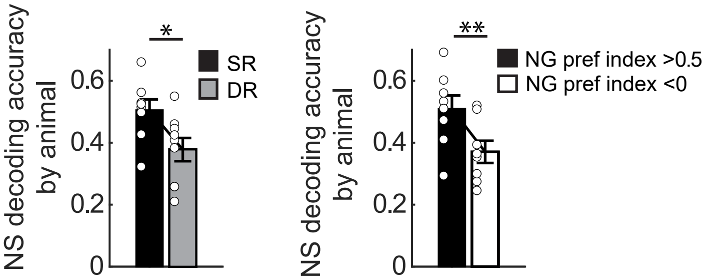


**Figure S4. Decoding results are robust to choice of classifier, related to Figures 3 and 5.**
Left, Decoding accuracy of natural scene (NS) stimuli was significantly decreased in dark-reared animals (DR, n= 9 animals) compared to standard-reared animals (SR, n= 8 animals; t-test, p= 0.026) using a Naive Bayes classifier. 151 neurons per animal were used, 24 trials for each natural scene stimulus were included. Right, In SR animals (n= 8), grating-preferring neurons (NG preference index <0) were selected for decoding natural scenes and compared to natural scene-preferring neurons (NG preference index > 0.5). Decoding accuracy was significantly lower in the grating-preferring group (paired t-test, p= 5.7e-4). The number of neurons was matched across the two groups on an animal-by-animal basis (neuron number by animal: 127,154, 106, 181, 139, 147, 108, 118).
Error bars indicate S.E.M. *p<0.05, **p<0.01

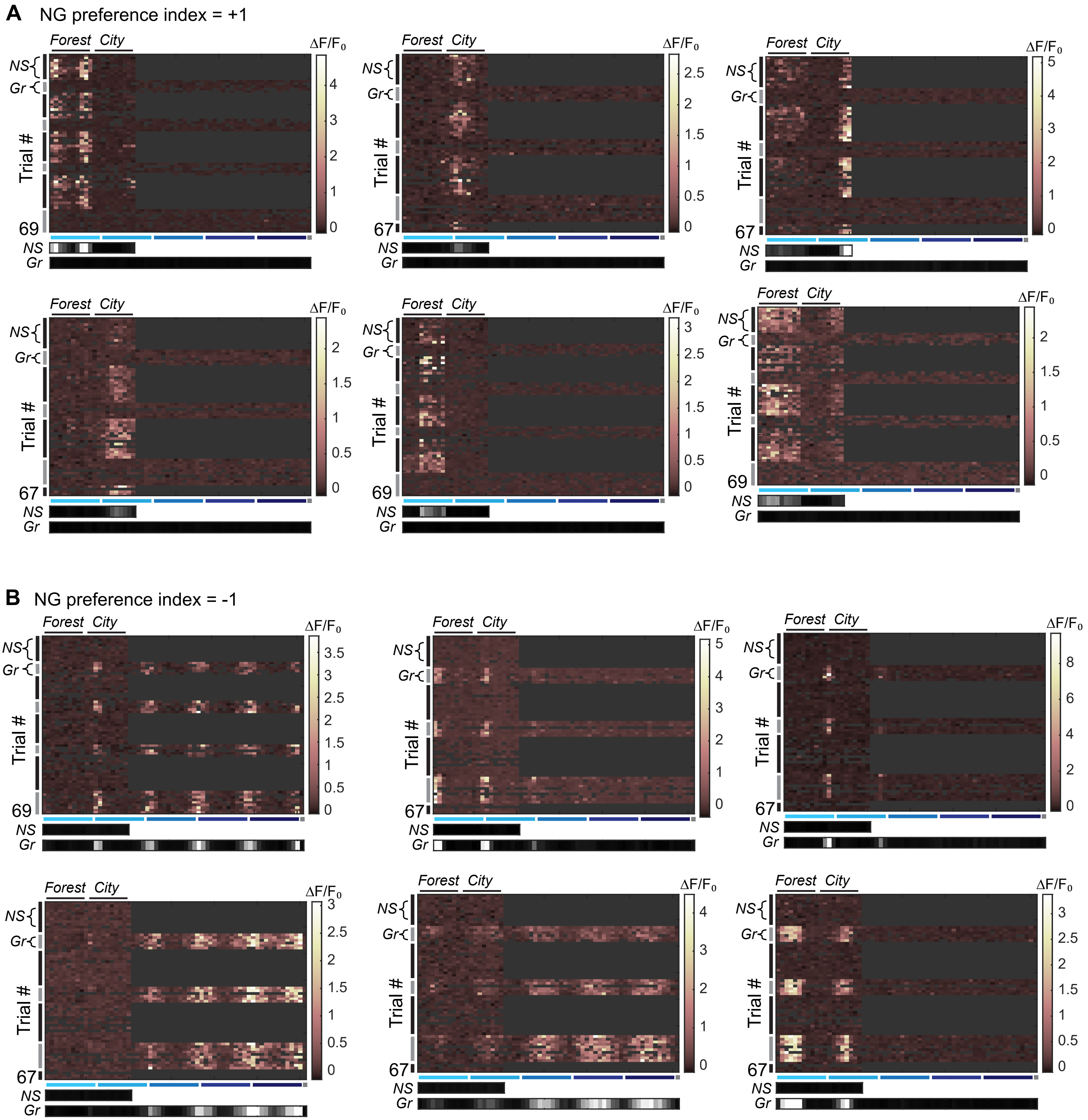


**Figure S5. Six example neurons that were not responsive to grating stimuli and six example neurons that were not responsive to natural scene stimuli, related to Figure 4.**
**A-B** Responses to grating and natural scene stimuli, Natural Scene (NS, black vertical bar) and grating stimuli (Gr, gray vertical bar) were presented in alternating blocks as indicated. The mean across NS and Gr trials is shown in gray scale at the bottom (scale limits: 1.5, 0 ΔF/F0). Stimulus IDs sorted as in Figure 1D,E.

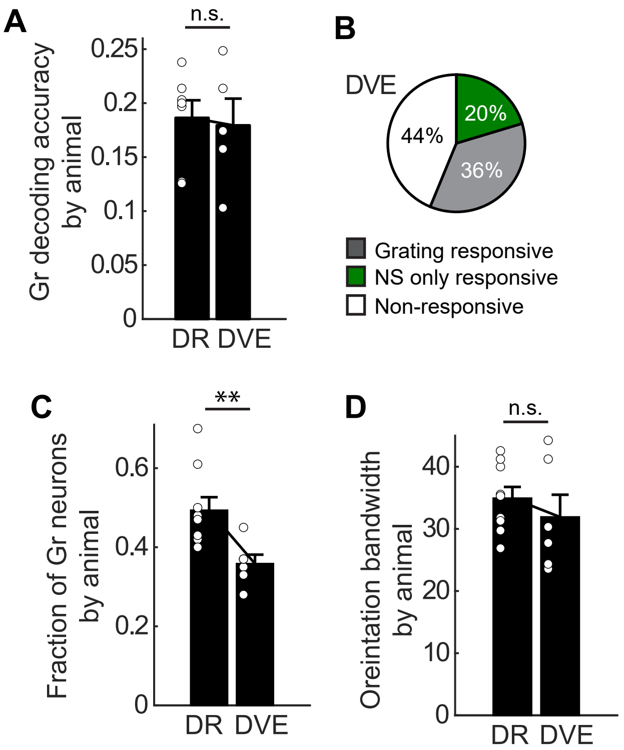


**Figure S6. Delayed visual experience reduced grating responsiveness but did not improve natural scene discriminability, realted to Figures 4 and 7.**

**A** Decoding accuracy of grating stimuli was not different between DR and DVE (n= 5 animals) animals (t-test p= 0.811). Only animals with at least 11 trials for each grating stimulus were included. 151 neurons per animal were used. Data from the DR condition were re-plotted from Fig. 7A.

**B** Proportion of neurons responding to stimuli as indicated, pooled across DVE animals (1633 total neurons from 6 animals). 49% of the grating responsive neurons also responded to at least one natural scene.

**C** The number of neurons responding to grating stimuli was significantly reduced in DVE animals (n= 6 animals) compared to DR animals (DR data was re-plotted from Fig. 2B; t-test, p= 4.8e-3).

**D** Median orientation bandwidth across animals was not different between DR and DVE (n= 6 animals) animals (t-test p= 0.423). Data from the DR condition was re-plotted from Fig. 7B.

Error bars indicate S.E.M. *p<0.05, **p<0.01

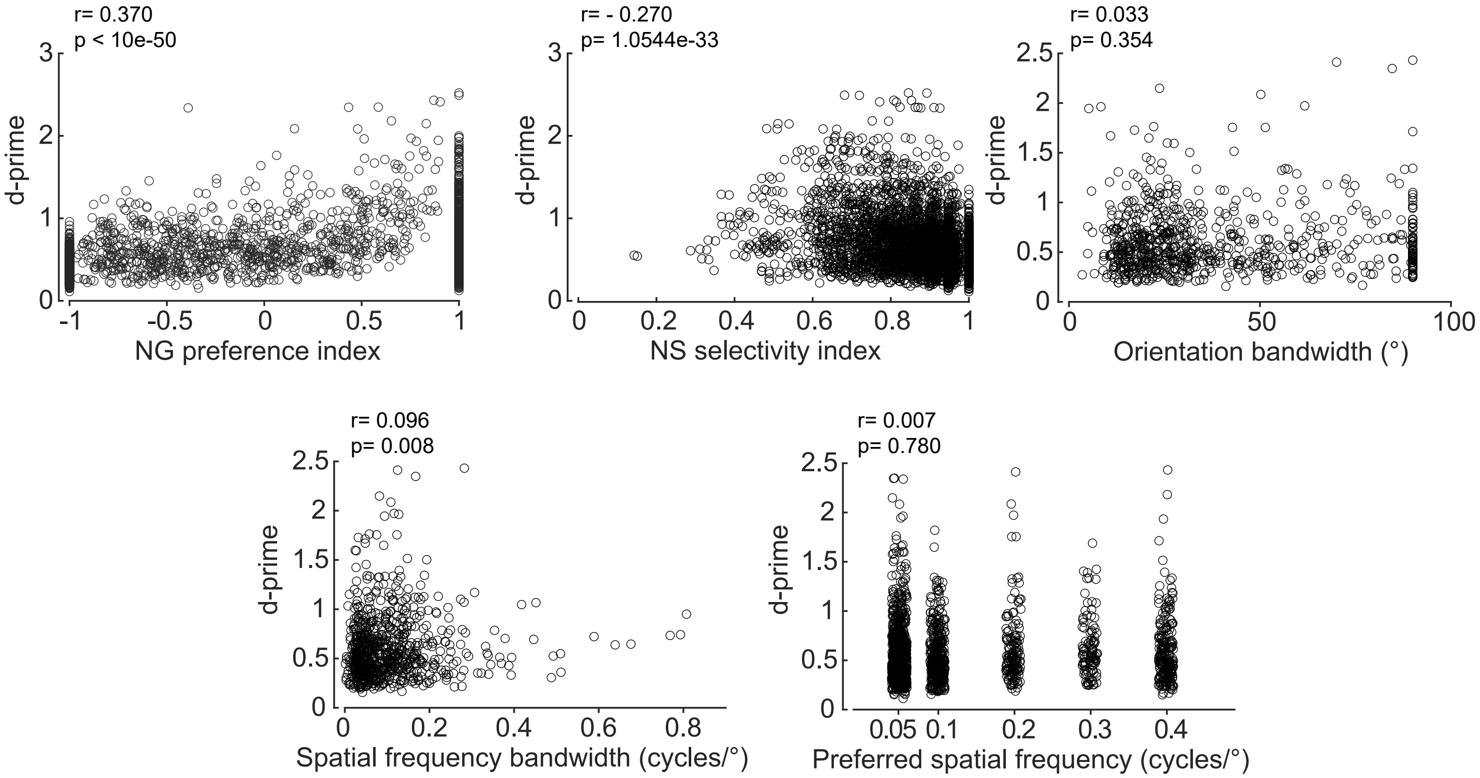


**Figure S7. NG preference index was positively correlated with natural scene d-prime, related to Figures 4 and 7.** Correlation (Spearman's rank) between natural scene d-prime and the five response properties as indicated. Note, the NS selectivity index was negatively correlated with natural scene d-prime. Visual inspection of the data points indicates that selectivity indices in the range of 0.65 to 0.85 have the highest d-prime values, and that the majority of selectivity indices above 0.9 have low d-prime values. These low d-prime values likely drive the negative correlation. Low d-prime values can be the result of trial-to-trial variability. Although the spatial frequency bandwidth p value was significant, the correlation value was ≤ 0.1, therefore we consider spatial frequency bandwidth to be uncorrelated with natural scene d-prime.

| Mouse ID | Sex | Doxycycline-treated | Age | # of responsive neurons |
|---|---|---|---|---|
| SR-1 | M | yes | 55 | 262 |
| SR-2 | M | yes | 51 | 296 |
| SR-3 | M | yes | 91 | 374 |
| SR-4 | F | yes | 96 | 418 |
| SR-5 | M | yes | 228 | 344 |
| SR-6 | F | yes | 168 | 302 |
| SR-7 | M | yes | 101 | 358 |
| SR-8 | M | yes | 54 | 257 |
| SR-9 | F | no | 28 | 172 |
| | | | 42 | 114 |
| SR-10 | M | no | 28 | 196 |
| | | | 44 | 136 |
| SR-11 | M | no | 29 | 250 |
| | | | 43 | 252 |
| | | | 81 | 181 |
| SR-12 | F | NA | 28 | 245 |
| | | | 42 | 227 |
| | | | 58 | 221 |
| SR-13 | M | NA | 28 | 258 |
| | | | 42 | 320 |
| | | | 59 | 338 |
| DR-1 | M | yes | 36 | 356 |
| | | | 89 (DVE) | 303 |
| DR-2 | F | yes | 37 | 497 |
| | | | 87 (DVE) | 247 |
| DR-3 | F | no | 38 | 271 |
| DR-4 | M | no | 40 | 380 |
| | | | 79 (DVE) | 314 |
| DR-5 | M | yes | 52 | 151 |
| DR-6 | M | yes | 46 | 262 |
| DR-7 | F | yes | 48 | 256 |
| DR-8 | M | yes | 43 | 298 |
| | | | 53 (DVE) | 335 |
| DR-9 | M | yes | 48 | 255 |
| | | | 64 (DVE) | 170 |
| DR-10 | F | yes | 89 (DVE) | 264 |

**Table S1. Animal information at the time of recording, related to Methods.**
The sex, age, and number of responsive neurons for each animal is indicated. Unless noted, mice are of the genotype: EMX1cre::Ai93D::CaMK2a-tTA. Five of the ‘DR’ animals listed here were transferred to a light cycle as adults, and the age of the second imaging session after light exposure is indicated. One DVE animal (DR-10) had a failure in eye tracking during the ‘DR’ imaging session. NA, not applicable, mice were of the genotype: SLC17a7cre:: Ai93D::CaMK2a-tTA.
